# Small molecule cognitive enhancer reverses age-related memory decline in mice

**DOI:** 10.1101/2020.04.13.039677

**Authors:** Karen Krukowski, Amber Nolan, Elma S. Frias, Morgane Boone, Gonzalo Ureta, Katherine Grue, Maria-Serena Paladini, Edward Elizarraras, Luz Delgado, Sebastian Bernales, Peter Walter, Susanna Rosi

## Abstract

With increased life expectancy age-associated cognitive decline becomes a growing concern, even in the absence of recognizable neurodegenerative disease. The integrated stress response (ISR) is activated during aging and contributes to age-related brain phenotypes. We demonstrate that treatment with the drug-like small-molecule ISR inhibitor ISRIB reverses ISR activation in the brain, as indicated by decreased levels of activating transcription factor 4 (ATF4) and phosphorylated eukaryotic translation initiation factor eIF2. Furthermore, ISRIB treatment reverses spatial memory deficits and ameliorates working memory in old mice. At the cellular level in the hippocampus, ISR inhibition i) rescues intrinsic neuronal electrophysiological properties, ii) restores spine density and iii) reduces immune profiles, specifically interferon and T cell-mediated responses. Thus, pharmacological interference with the ISR emerges as a promising intervention strategy for combating age-related cognitive decline in otherwise healthy individuals.

**ONE SENTENCE SUMMARY:** Inhibition of the integrated stress response restores neuronal and immune dysfunction and alleviates memory deficits in aged mice.

## INTRODUCTION

“Of the capacities that people hope will remain intact as they get older, perhaps the most treasured is to stay mentally sharp” (*1*).

The impact of age on cognitive performance represents an important quality-of-life and societal concern, especially given our prolonged life expectancy. While often discussed in the context of disease, decreases in executive function as well as learning and memory decrements in older, healthy individuals are common (*2*, *3*, *4*, *5*). According to the US Department of Commerce the aging population is estimated by 2050 to reach 83.7 million individuals above 65 years of age in the US; this represents a rapidly growing healthcare and economic concern (*6*).

Age-related decline in memory has been recapitulated in preclinical studies with old rodents (*7*–*10*). Specifically, prior studies have identified deficits in spatial memory (*9*, *11*), working and episodic memory (*8*, *10*) and recognition memory (*12*), when comparing young, adult mice with older sex-matched animals. The hippocampus is the brain region associated with learning and memory formation and is particularly vulnerable to age-related changes in humans and rodents (*13*–*16*). Deficits in a number of cellular processes have been suggested as underlying causes based on correlative evidence, including protein synthesis (*17*), metabolism (*18*), inflammation (*19*), and immune responses (*9*, *11*, *20*, *21*). While providing a wealth of parameters to assess, by and large the causal molecular underpinnings of age-related memory decline have remained unclear.

The principle that blocking protein synthesis prevents long-term memory storage was discovered many years ago (*22*). With age there is a marked decline of protein synthesis in the brain that correlates with defects in proper protein folding (*12*, *23*–*25*). Accumulation of misfolded proteins can activate the integrated stress response (ISR) (*26*), an evolutionary conserved pathway that decreases protein synthesis. In this way, the ISR may have a causative role in age-related cognitive decline. We previously discovered that interference with the drug-like small-molecule inhibitor (integrated stress response inhibitor, or ISRIB) rescued traumatic brain injury-induced behavioral and cognitive deficits (*27*–*29*), suggesting that this pharmacological tool may be useful in testing this notion.

Increasing age leads to structural and functional changes in hippocampal neurons. Specifically, in old animals there is an increase in neuronal hyperpolarization after spiking activity (“afterhyperpolarization”, or AHP) that decreases intrinsic neuronal excitability and correlates with memory deficits (*13*–*16*, *30*). Aging also manifests itself with synaptic excitability changes in the hippocampus that correlate with a reduction in the bulbous membrane projections that form the postsynaptic specializations of excitatory synapses, termed dendritic spines (*31*, *32*). Morphological changes in dendritic spine density are critical for spatial learning and memory (*33*, *34*). Whether these age-related neuronal changes can be modified or are linked with ISR activation has yet to be determined.

In addition to neuronal changes, ISR activation can modify immune responses via alterations in cytokine production (*35*). Indeed, maladaptive immune responses have been linked with cognitive decline in the old brain (*8*, *9*, *11*, *20*). Initial studies focused on age-associated cytokine responses, including interferon (IFN)-mediated cognitive changes (*20*, *36*). Type-I IFN responses can induce age-related phenotypes in rodents. Furthermore, the adaptive immune system (T cell infiltration into the old brain) can regulate neuronal function via IFN-γ production (*21*), suggesting the possibility that age-induced maladaptive immune responses and the ISR are linked. Here we explore the possibility of ISR inhibition by ISRIB as a potential strategy for modifying age-induced neuronal, immune, and cognitive dysfunction.

## RESULTS

### ISRIB resets the ISR in the brain of old mice

ISR activation leads to global reduction in protein synthesis but also to translational up-regulation of a select subset of mRNAs whose translation is controlled by small upstream open-reading frames in their 5’-UTRs (*37*, *38*). One well-studied ISR-upregulated target protein is ATF4 (activating transcription factor 4) (*39*, *40*). We recently showed ISRIB administration reversed mild head trauma-induced elevation in ATF4 protein (*28*). Using the same ISRIB treatment paradigm of daily injections on 3 consecutive days (*27*, *28*), we found decreased age-associated ATF4 protein levels in mouse brain lysates when compared to vehicle-treated controls during ISRIB administration (**Supplemental Figure 1**). ATF4 levels 18 days after cessation of ISRIB treatment showed persistent reduction in age-induced ATF4 protein levels that were indistinguishable from young mice (**Figure 1A, B**, **Supplemental Figure 2A**).

**Figure 1.**
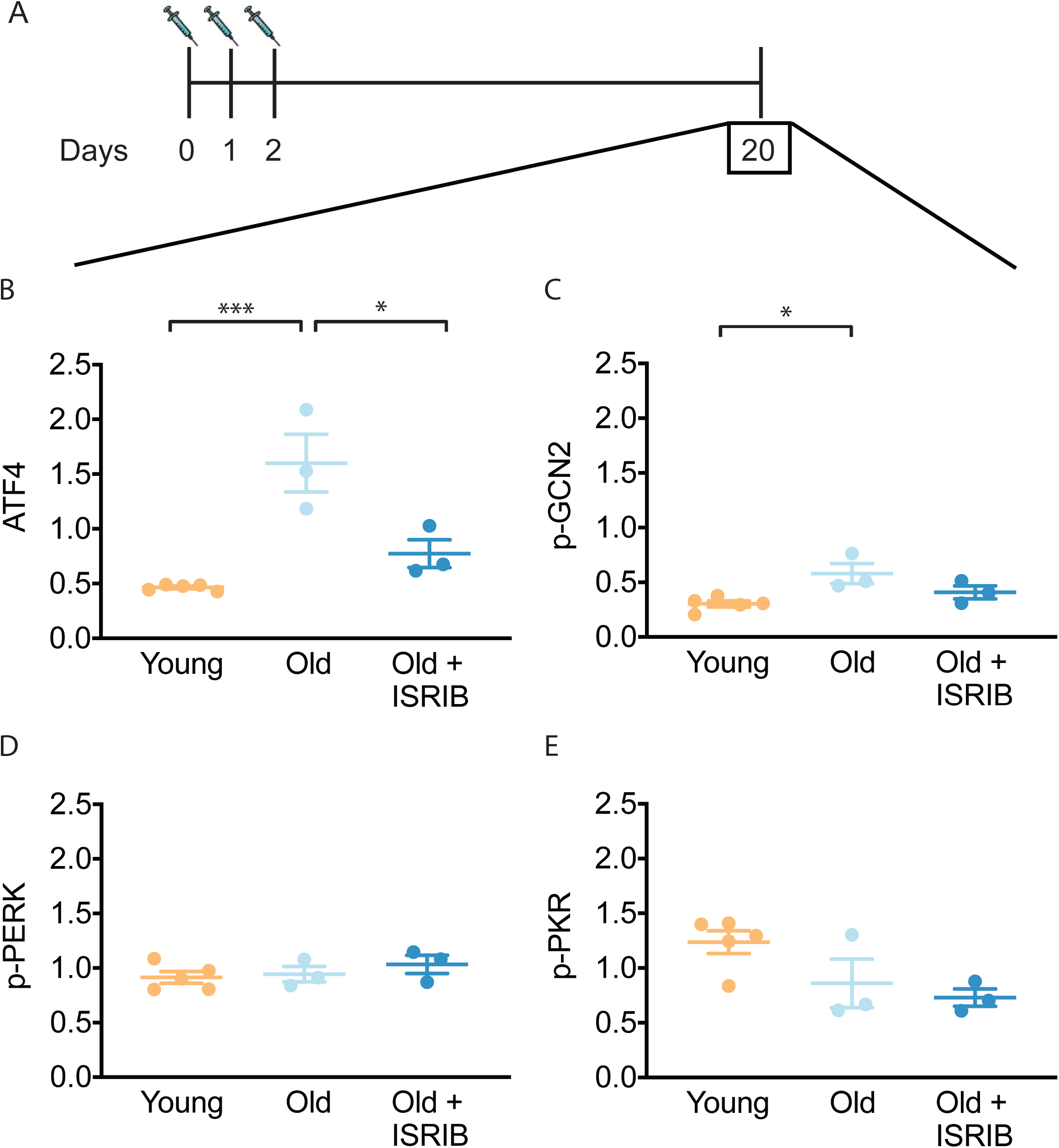
ISRIB resets the ISR in the brain of old mice. (**A**) Experimental dosing scheme: ISRIB treatment denoted by syringes (3 injections). (**B**) ISRIB treatment reduced ATF4 protein levels chronically 18 days after ISRIB treatment was complete. One-way ANOVA (F = 18.8, p < 0.001); with Tukey post-hoc analysis. (**C**) Modest age-induced increases in p-GCN2 when comparing young and old male mice. One-way ANOVA (F = 6.6, p < 0.05); with Tukey post-hoc analysis. (**D**, **E**) Age and ISRIB administration did not impact p-PERK or p-PKR protein levels. Brain lysates of specific protein levels listed normalized to actin. Young n = 5, Old = 3, Old + ISRIB = 3. Individual animal values represented by dots; lines depict group mean ± SEM. *p < 0.05; ***p < 0.001

The key regulatory step in the ISR lies in the phosphorylation of eukaryotic translation initiation factor eIF2 (*26*). Four known kinases can phosphorylate Ser51 in its α-subunit of (eIF2α) to activate the ISR (*41*): HRI (heme-regulated inhibitor), PKR (double-stranded RNA-dependent protein kinase), PERK (PKR-like ER kinase) and GCN2 (General amino acid control nonderepressible 2). Only three of these kinases are known to be expressed in the mammalian brain (PKR, PERK, GCN2). To understand upstream modifiers of age-related ISR activation, we investigated the impact of age and ISRIB administration on the expression of these kinases. We found a modest, but significant increase in activated GCN2 (as indicated by its phosphorylated form p-GCN2) when comparing young and old brain lysates (**Figure 1C**, **Supplemental Figure 2B**). Moreover, when ISRIB was administered weeks prior (**Figure 1A**), GCN2 activation returned to levels comparable to young brains (**Figure 1C**, **Supplemental Figure 2B**). Age and ISRIB did not impact phosphorylation status of PERK and PKR in total brain lysates (**Figure 1D, E**, **Supplemental Figure 2B**). Thus, brief ISRIB administration in the old brain has long-lasting effects on ISR activation.

### Inhibition of the ISR reverses age-induced decline in spatial learning and memory

To assess whether the reduction in ISR activation affects age-related cognitive defects, we tested the capacity for spatial learning and memory in young and old mice in a radial arm water maze (*27*, *42*). This particular forced-swim behavior tool measures hippocampal-dependent spatial memory functions in rodents and has been previously used to assess age-related cognitive deficits (*7*, *43*). Animals were trained for two days (two learning blocks/day) to locate a platform hidden under opaque water in one of the eight arms using navigational cues set in the room (**Figure 2A**). We recorded the total number of entries into the non-target arm (errors) before the animal found the escape platform with automated tracking software and used it as a metric of learning. After two days of training, young animals averaged one error prior to successfully locating the escape platform, while old animals averaged three errors, indicating their reduced learning capacity (**Supplemental Figure 3A**).

**Figure 2.**
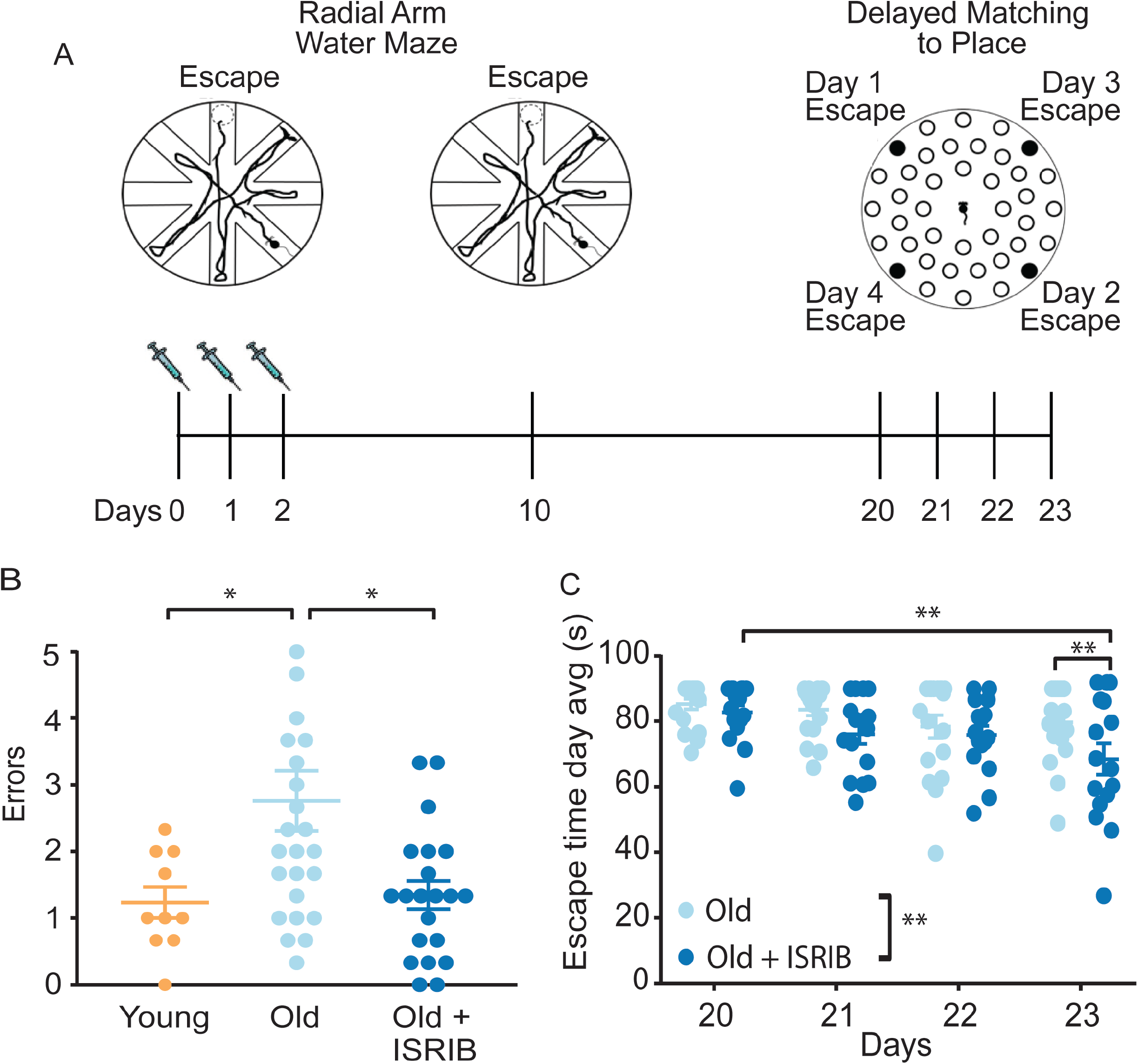
Inhibition of the ISR reverses age-induced decline in spatial, working and episodic memory. (**A**) Experimental Design: Old (~19 months) animals underwent behavioral analysis in a radial arm water maze (RAWM) and a delayed matching to place paradigm (DMP). ISRIB or vehicle administration (2.5 mg/kg intraperitoneal) occurred daily during the learning phase of RAWM denoted by syringes (days 0-2). (**B**) ISRIB treatment improved memory one week after administration in male rodents. One-way ANOVA (F = 5.3, p < 0.05); with Tukey post-hoc analysis. Young n = 10; Old n = 25; Old + ISRIB n = 21. (**C**) Age-induced deficits in working and episodic learning and memory restored weeks after ISRIB administration. Animals performed the DMP from day 20 – 23. Average of all trials per group for each day. Day 20, 21 = 4 trials/day. Day 22,23 = 3 trials/day. Two-way repeated measures ANOVA reveals a significant difference between groups p < 0.01 (denoted in figure legend) and time effect p < 0.01. *p < 0.05, **p < 0.01. Old n = 18; Old + ISRIB n = 16. Individual animal scores represented by dots; lines depict group mean ± SEM.

We next tested whether pharmacological inhibition of the ISR could modify the age-related spatial learning deficits. ISRIB treatment started the day before the first training day and continued with daily injections throughout the duration of the training (3 injections in total; see **Figure 2A**, left). By the end of two days of training, ISRIB-treated old animals averaged two errors prior to finding the escape platform, while vehicle-treated old animals averaged three, denoting significant learning improvement in the mice that received ISRIB (**Supplemental Figure 3B**). No difference in learning performance was measured in young mice that received the identical treatment paradigm (**Supplemental Figure 3C**), suggesting that ISRIB-induced learning improvement measured in this training regime is age-dependent. These results were confirmed in an independent old animal cohort, in which we tested an additional ISR inhibitor (Cmp-003, a small molecule with improved solubility and pharmacological properties (PCT/US18/65555)), using an identical training/injection paradigm (**Supplemental Figure 3D**). Old animals that received Cmp-003 made significantly fewer errors prior to locating the escape platform than old animals that received vehicle injections, again indicating significant learning improvement.

Spatial memory of the escape location was measured one week later by reintroducing the animals into the pool and measuring the number of errors before they located the hidden platform. The animals did not receive any additional treatment during this task. Old mice treated with ISRIB one week before made significantly fewer errors compared to matched, vehicle-treated old male (**Figure 2B**) and female (**Supplemental Figure 4**) mice. Remarkably, the memory performance of old animals treated with ISRIB a week before was comparable to that of young mice (**Figure 2B**). These results demonstrate that brief treatment with ISRIB rescues age-induced spatial learning and memory deficits, cementing a causative role of the ISR on long-term memory dysfunction.

### ISRIB administration improves age-induced deficits in working and episodic memory weeks after treatment

Given the long-lasting effect of brief ISRIB treatment on ATF4 protein levels in the brain and on memory function one week after drug administration, we next tested the duration of ISRIB effects on age-related cognitive function. On experimental day 20 (18 days post ISRIB treatment, **Figure 2A**, right), we measured working and episodic memory using a delayed-matching-to-place paradigm (DMP) (*27*, *44*) in the same animal cohort without additional ISRIB treatment. Previous work has demonstrated that old mice display significant impairments when compared to young mice (*8*, *10*). During DMP animals learned to locate an escape tunnel attached to one of 40 holes in a circular table using visual cues. The escape location was changed daily, forcing the animal to relearn its location. To quantify performance, we used analysis tracking software to measure “escape latency”, reporting the time taken by the mouse to enter the escape tunnel.

Old mice that received ISRIB treatment 18 days earlier displayed significant improvement over the four-day testing period (**Figure 2C**; Day 20 vs. Day 23). By Day 23 post-treatment animals were locating the escape tunnel on average 20 seconds faster than the matched-vehicle group (**Figure 2C**). This behavior is indicative of improved working and episodic memory. By contrast, old animals that received vehicle injections did not learn the task (**Figure 2C**; Day 20 vs. Day 23), as previously observed (*8*, *10*). These results demonstrate that ISRIB administration increases cognitive performance in a behavioral paradigm measured weeks after administration.

### ISRIB treatment reverses age-associated changes in hippocampal neuron function

To determine the neurophysiological correlates of ISRIB treatment on cognition, we investigated its effects on hippocampal neuronal function. Utilizing whole cell patch clamping, we recorded intrinsic electrophysiological firing properties and synaptic input in CA1 pyramidal neurons of young and old mice and compared them to those of old mice treated with a single injection of ISRIB the day prior to recording (**Figure 3A**).

**Figure 3.**
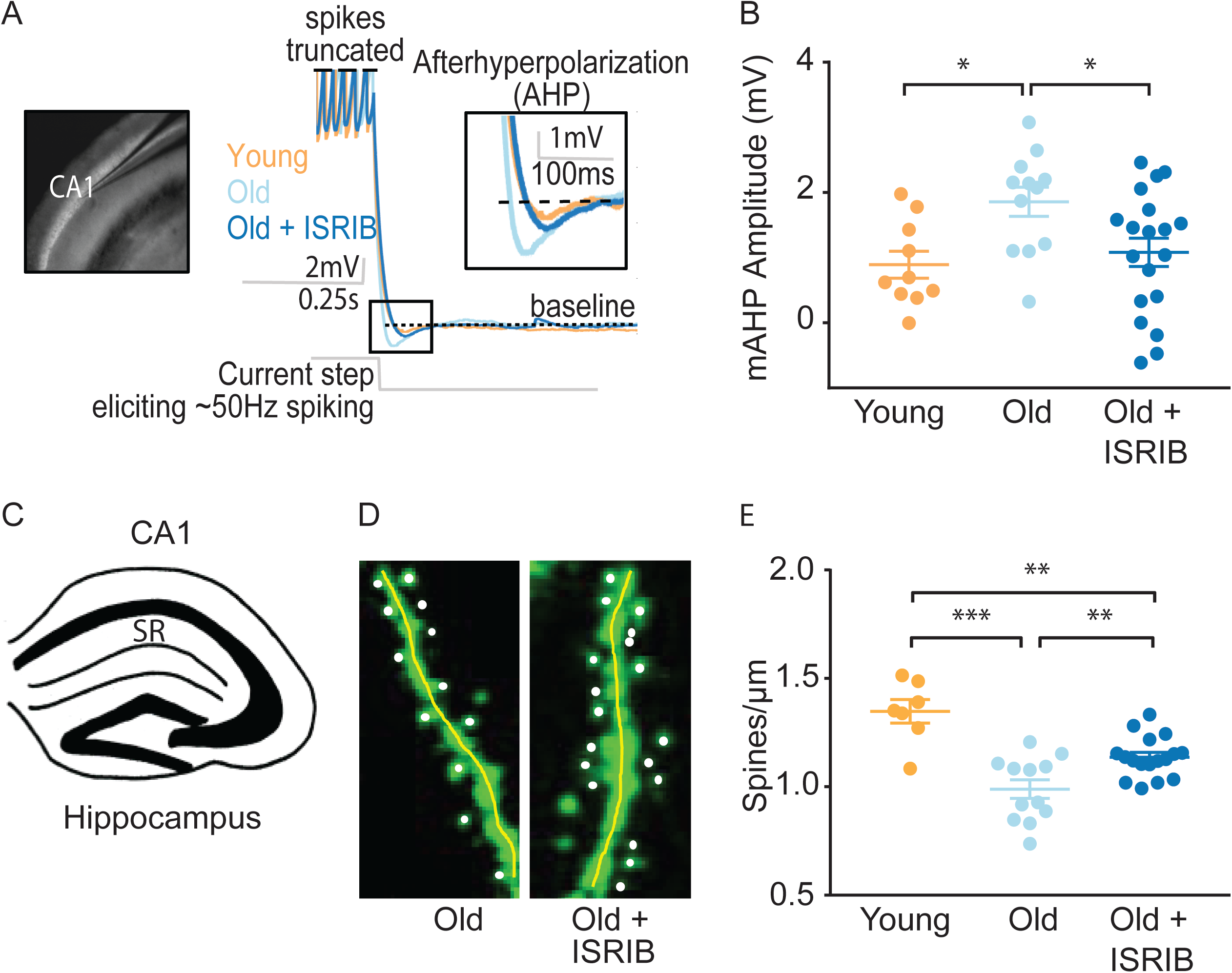
ISRIB treatment alleviates age-associated changes in CA1 pyramidal neuron function and structure. (**A**) Left: Image of pipette patched onto CA1 neuron in sagittal slice of hippocampus. Right: Representative traces from hippocampal CA1 pyramidal neurons from old animals treated with either vehicle (light blue) or ISRIB (dark blue) or young animals treated with vehicle (orange) showing the response to a current injection eliciting ~50Hz spiking activity. Spikes are truncated (dashed line), and the AHP is visualized immediately following cessation of current injection (yellow square) and quantified as the change in voltage from baseline (dotted line). (**B**) Age-induced increases in AHP were measured when comparing young and old animals. ISRIB treatment reversed increased AHP to levels indistinguishable from young animals. Animals were injected with ISRIB (2.5 mg/kg) or vehicle intraperitoneal one day prior to recordings. One-way ANOVA (F = 4.461, p < 0.05); with Tukey post-hoc analysis. *p < 0.05. Each neuron is represented with a symbol; lines indicate the mean ± SEM (Neurons: Young males n = 10 (5 animals); Old males n = 12 (5 animals), Old + ISRIB males n = 19 (7 animals) with 1-5 neurons recorded per animal. (**C-E**) Spine density was quantified in the CA1 region of the dorsal hippocampus from young and old Thy1-YFP-H mice. (**C**) Diagram of hippocampal region analyzed. SR = stratum radiatum. (**D**) Representative images from Old and Old + ISRIB mice. (**E**) A decrease in dendritic spine density was measured when comparing old mice to young mice. ISRIB treatment significantly increased spine density levels of old mice when compared to vehicle treated old mice. 63x magnification with a water immersion objective. Young males n = 7 slides (2 animals); Old males + Vehicle n = 12 slides (3 mice); Old males + ISRIB n = 17 slides (4 mice). Individual slide scores (relative to old mice) represented in dots, lines depict group mean ± SEM. One-way ANOVA (F = 18.57, p < 0.001) with Tukey post-hoc analysis. **p < 0.01; ***p < 0.001.

We evaluated alterations in intrinsic excitability by measuring action potential shape and frequency properties and passive membrane response properties produced by a series of hyperpolarizing and depolarizing current steps (20 steps from −250 to 700 pA, 250 ms duration). We also assessed the hyperpolarization of the membrane potential following high frequency firing, specifically the AHP following ~50 Hz spiking activity induced with a current step (**Figure 3A**). In agreement with previous reports (*13*–*16*, *30*), old mice displayed a significantly increased AHP amplitude when compared to young mice (**Figure 3B**). ISRIB treatment reversed the age-induced increase in AHP amplitude, rendering the CA1 neuronal response in ISRIB-treated old mice indistinguishable from young mice (**Figure 3B**). We did not find significant differences in other action potential or passive membrane properties between groups (**Supplemental Figure 5**).

We also measured spontaneous excitatory postsynaptic currents (sEPSC), while holding the cell at −75 mV in a voltage clamp. Both the frequency and amplitude of sEPSCs were indistinguishable between groups (**Supplemental Figure 6**). These data support that ISRIB treatment in old animals restores neuronal function to levels comparable to young neurons by affecting intrinsic excitability and specifically reducing the AHP following high frequency firing.

### ISRIB treatment reduces dendritic spine loss

To determine if ISRIB might affect age-induced synaptic structural changes, we quantified dendritic spine density after ISRIB treatment in old mice with fluorescently labeled excitatory neurons (marked by a genomically encoded Thy1-YFP fusion protein). The hippocampus of old mice is characterized by a reduction in dendritic spine density that correlates with diminished cognitive output (*31*, *32*). Old Thy1-YFP expressing mice received ISRIB treatment and two days of behavioral training as described in **Figure 2A**. At the end of Day 2, we terminated the animals and harvested the brains for quantification of dendritic spine density in the hippocampus (stratum radiatum of CA1) (**Figure 3C**) using confocal microscopy imaging and unbiased analysis (**Figure 3D**). Similar to previous reports, we measured a significant reduction in dendritic spine density in old when compared to young Thy1-YFP mice (**Figure 3E**) (*31*, *32*). ISRIB treatment significantly increased spine numbers when compared with age-matched vehicle-treated mice (**Figure 3E**). Taken together these data demonstrate that ISRIB administration improves both neuron structure and function in old mice.

### Age-induced inflammatory tone is reduced following ISRIB treatment

Because it is known that immune dysregulation in the brain increases with age (*45*) and correlates with reduced cognitive performance in old animals (*8*, *9*, *11*, *20*), we next investigated immune parameters impacted by ISRIB administration in the old brain. To this end, we first investigated glial cell activation (microglia and astrocytes) in hippocampal sections from of old mice and ISRIB-treated old mice by fluorescent microscopic imaging (during ISRIB administration, **Supplemental Figure 7**). We measured, astrocyte and microglia reactivity as the percent area covered by GFAP and Iba-1. In these analyses, we observed no differences between the old and ISRIB-treated old animals (**Supplemental Figure 7B-G**).

Next, we performed quantitative PCR (qPCR) analyses on hippocampi from young, old, and ISRIB-treated animals on samples taken at the same time point as in the microscopic analysis (**Supplemental Figure 7A**). We measured a panel of inflammatory markers, many of which are known to increase with age (*45*), with a particular focus on IFN-related genes as this pathway is implicated in age-related cognitive decline (*20*, *36*). Indeed, we found that age increased expression of a number of IFN response pathway genes, *Rtp4, Ifit1, and Gbp10* (**Figure 4A-C**). Importantly, ISRIB administration reduced expression of *Rtp4, Ifit1, and Gbp10* to levels that became indistinguishable from young animals (**Figure 4A-C**). Other inflammatory makers were also increased with age (*Ccl2, Il6*) but not affected by ISRIB treatment, whereas *Cd11b* was increased upon ISRIB administration alone (**Table 1**).

**Figure 4.**
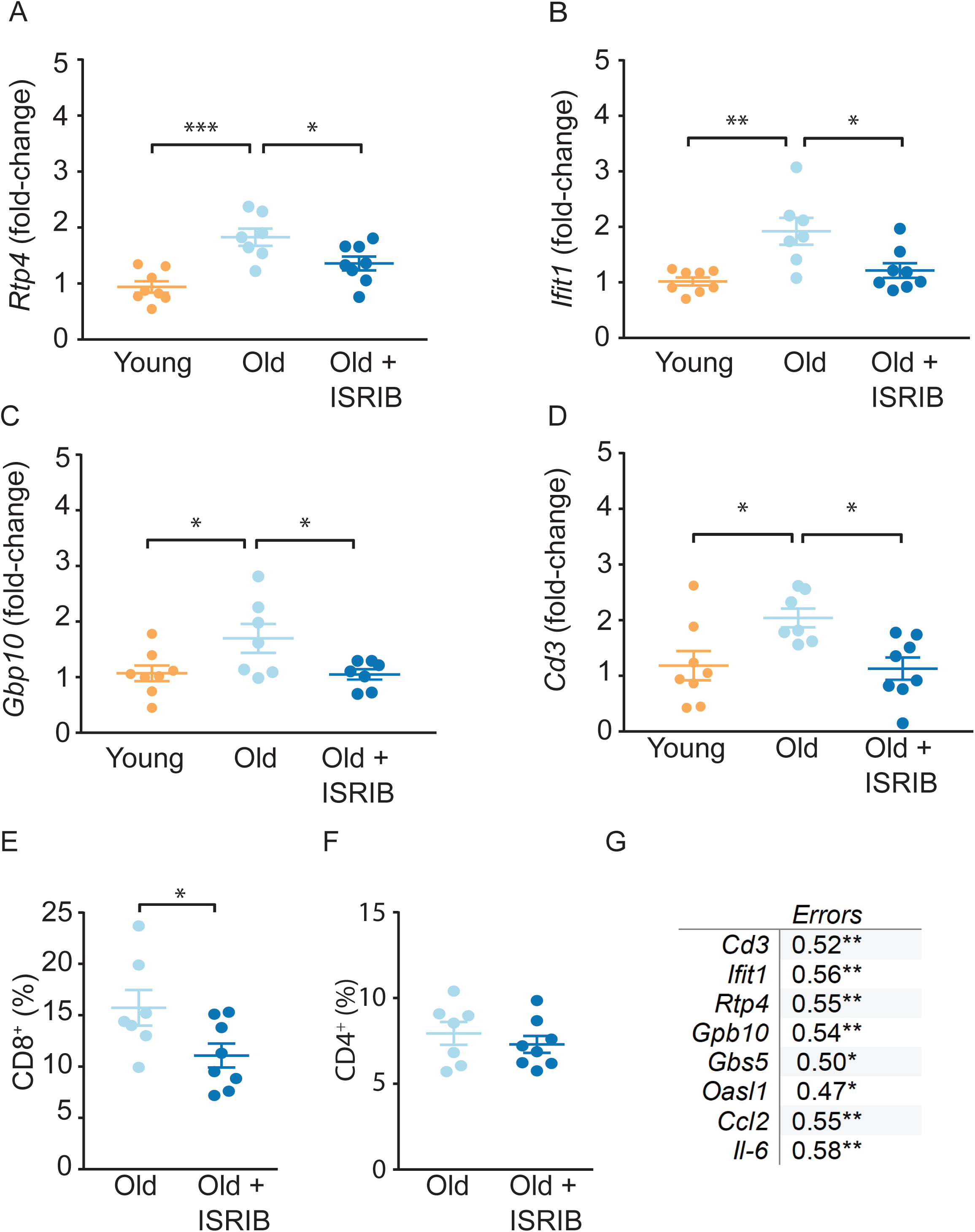
Age-induced inflammatory tone is reduced following ISRIB treatment. Inflammatory genes were investigated in the hippocampus of young and old mice by qPCR analysis. (**A**-**C**) ISRIB administration reversed age-induced increases in *Rtp4*, *Ifit1*, and *Gbp10*. (**A**) *Rtp4*, One-way ANOVA (F = 12.23, p < 0.001) with a Tukey-post analysis. Young males n = 8; Old males n = 7; Old + ISRIB males n = 8. (**B**) *Ifit1*, One-way ANOVA (F = 8.8; p < 0.01) with a Tukey-post analysis. Young males n = 8; Old males n = 7; Old + ISRIB males n = 8. (**C**) *Gbp10*, One-way ANOVA (F =4.2, p < 0.05) with a Tukey-post analysis. Young males n = 8; Old males n = 7; Old + ISRIB males n = 7. (**D**) *Cd3* gene-expression (a marker for T cells) changes in the hippocampus of young and old animals were measured by qPCR analysis. *Cd3* was significantly increased with age. ISRIB administration returned *Cd3* expression levels to those comparable to young animals. One-way ANOVA (F = 5.2; p < 0.05). Tukey-post hoc analysis. Young males n = 8; Old males n = 7; Old + ISRIB males n = 8. (**E**, **F**) Peripheral T cell levels were measured by flow cytometric analysis of whole blood. (**E**) ISRIB treatment reduced CD8+ T cell percentages (of CD45+ cells) in the peripheral blood. Student t-test. Old males n = 7; Old + ISRIB males n = 8. (**F**) CD4+ T cell percentages (of CD45+ cells) were not impacted. Individual animal scores represented by dots; lines depict group mean ± SEM. (**G**) A significant positive correlation was measured between cognitive performance on day 2 of the RAWM (errors) and multiple inflammatory markers (*Cd3*, *Ifit1*, *Rtp4*, *Gbp10*, *Gbp5*, *Oasl1*, *Ccl2*, *Il-6*). Linear regression was measured by Pearson R correlation, R value denoted with significance. *p < 0.05; **p < 0.01; ***p < 0.001.

**Table 1.** Impact of age and ISRIB on mRNA expression in the hippocampus. Inflammatory, ISR mediators and neuronal health targets were investigated by qPCR analysis of hippocampal lysates after two ISRIB injections. Columns: (i) mRNA targets (ii) Young group mean ± SEM (iii) Old group mean ± SEM (iv) Old + ISRIB group mean ± SEM (v) ANOVA F value (vi) Significant denotation between groups (vii) n/group.

| **TARGET** | **Young** | **Old** | **Old + ISRIB** | **ANOVA** | **Btw Groups** | **n: Yg/Old/Old + ISRIB** |
| --- | --- | --- | --- | --- | --- | --- |
| ***Ccl2*** | 1.0 ± 0.1 | 2.3 ± 0.4 | 1.6 ± 0.1 | F = 5.6* | Yg v Old** | 8/7/7 |
| ***Cd11b*** | 1.0 ± 0.0 | 1.0 ± 0.0 | 1.2 ± 0.0 | F = 6.6** | Yg v Old +<br>ISRIB**<br>Old v Old +<br>ISRIB* | 8/7/8 |
| ***Il1-β*** | 1.1 ± 0.1 | 1.4 ± 0.1 | 1.7 ± 0.1 | F = 3.6* | Yg v Old +<br>ISRIB* | 8/7/8 |
| ***Tnfα*** | 1.0 ± 0.1 | 1.6 ± 0.3 | 2.1 ± 0.4 | F = 2.7 |  | 7/7/8 |
| ***Il-6*** | 1.1 ± 0.2 | 1.8 ± 0.1 | 1.8 ± 0.2 | F = 4.1* | Yg v Old* | 8/7/8 |
| ***Il-10*** | 1.0 ± 0.1 | 1.4 ± 0.2 | 1.5 ± 0.2 | F = 1.1 |  | 7/7/7 |
| ***Irf7*** | 1.0 ± 0.1 | 1.4 ± 0.1 | 1.2 ± 0.1 | F = 1.5 |  | 8/7/8 |
| ***Ifitm3*** | 1.0 ± 0.0 | 1.2 ± 0.1 | 1.0 ± 0.0 | F = 0.8 |  | 8/7/8 |
| ***Isg15*** | 1.0 ± 0.0 | 1.3 ± 0.1 | 1.2 ± 0.0 | F = 2.0 |  | 8/7/7 |
| ***Ifi204*** | 1.1 ± 0.2 | 1.4 ± 0.2 | 1.5 ± 0.1 | F = 1.2 |  | 6/7/7 |
| ***Gbp5*** | 1.0 ± 0.0 | 1.2 ± 0.1 | 1.1 ± 0.0 | F = 2.6 |  | 8/7/8 |
| ***Oasl1*** | 1.0 ± 0.0 | 1.8 ± 0.3 | 0.9 ± 0.1 | F = 4.9* | Yg v Old*;<br>Old v Old +<br>ISRIB* | 8/7/8 |
| ***Ophn1*** | 1.0 ± 0.0 | 0.9 ± 0.0 | 1.1 ± 0.0 | F = 1.8 |  | 8/7/8 |
| ***Bdnf*** | 1.0 ± 0.0 | 0.9 ± 0.0 | 1.0 ± 0.0 | F = 2.7 |  | 8/7/8 |
| ***Gadd34*** | 1.0 ± 0.0 | 0.8 ± 0.0 | 0.9 ± 0.0 | F = 1.8 |  | 7/7/8 |
| ***Pkr*** | 1.0 ± 0.0 | 1.0 ± 0.0 | 1.1 ± 0.0 | F = 1.0 |  | 8/7/8 |

Using these same hippocampal extracts, we next measured T cell responses. Similar to other reports (*21*, *46*), we observed a significant increase in T cell marker mRNA expression (*Cd3*) in the hippocampus of old compared to young mice (**Figure 4D**). ISRIB treatment in the old mice reduced the expression of the T cell marker to a level comparable to that observed in young mice (**Figure 4D**). The ISRIB-induced reduction in T cell marker levels was not limited to the brain but extended to the peripheral blood of old animals, with CD8+ T cell percentages reduced following ISRIB administration (**Figure 4E**). By contrast, we observed no changes in CD4+ T cell levels (**Figure 4F**).

Given the broad and varied response of immune parameters in response to ISRIB treatment, we next explored possible relationships between behavioral performance and age-related inflammatory tone. T cell marker mRNA expression in the brain positively correlated with cognitive performance: mice with lower T cell marker expression made fewer errors prior to locating the escape platform during memory testing on Day 2 (**Figure 4G**). We also observed significant positive correlations when comparing memory performance on day 2 (errors) with the mRNA levels of multiple IFN response pathway genes (*Ifit1, Rtp4, Gbp10, Gbp5, Oasl1*) and additional inflammatory markers (*Ccl2, Il6*) (**Figure 4G**). These data demonstrate that increased inflammatory marker expression correspond with poorer cognitive performance, even when we did not observed differences between the groups. These studies revealed that ISRIB treatment impacts a broad number of immune parameters both in the periphery and in the brain reducing the age-related inflammatory tone which strongly correlates with improved cognition.

### ISRIB treatment resets age-related ISR activation

Finally, we investigated the transcriptional expression of ISR mediators and neuronal health markers in the hippocampal lysates used above. We did not detect differences in *Gadd34, Pkr, Bdnf1,* and *Ophn1* mRNA levels with age or ISRIB treatment (**Table 1**). Interestingly however, when analyzed as individual animals, we found a negative correlation between *Gadd34* mRNA and cognitive performance: animals with less *Gadd34* mRNA made more errors prior to locating the escape platform during memory testing (**Figure 5A**). GADD34, the regulatory subunit of one of the two eIF2α phosphatases, acts in a feedback loop as a downstream target of ATF4 (*47*–*49*). Induction of GADD34 leads to decreased phosphorylation of eIF2 which counteracts ISR activation. Indeed, ISRIB treatment reduced p-eIF2 levels in total brain lysates (**Figure 5B; Supplemental Figure 8**), suggesting that ISRIB may break a feedback loop thereby resetting age-related ISR activation.

**Figure 5.**
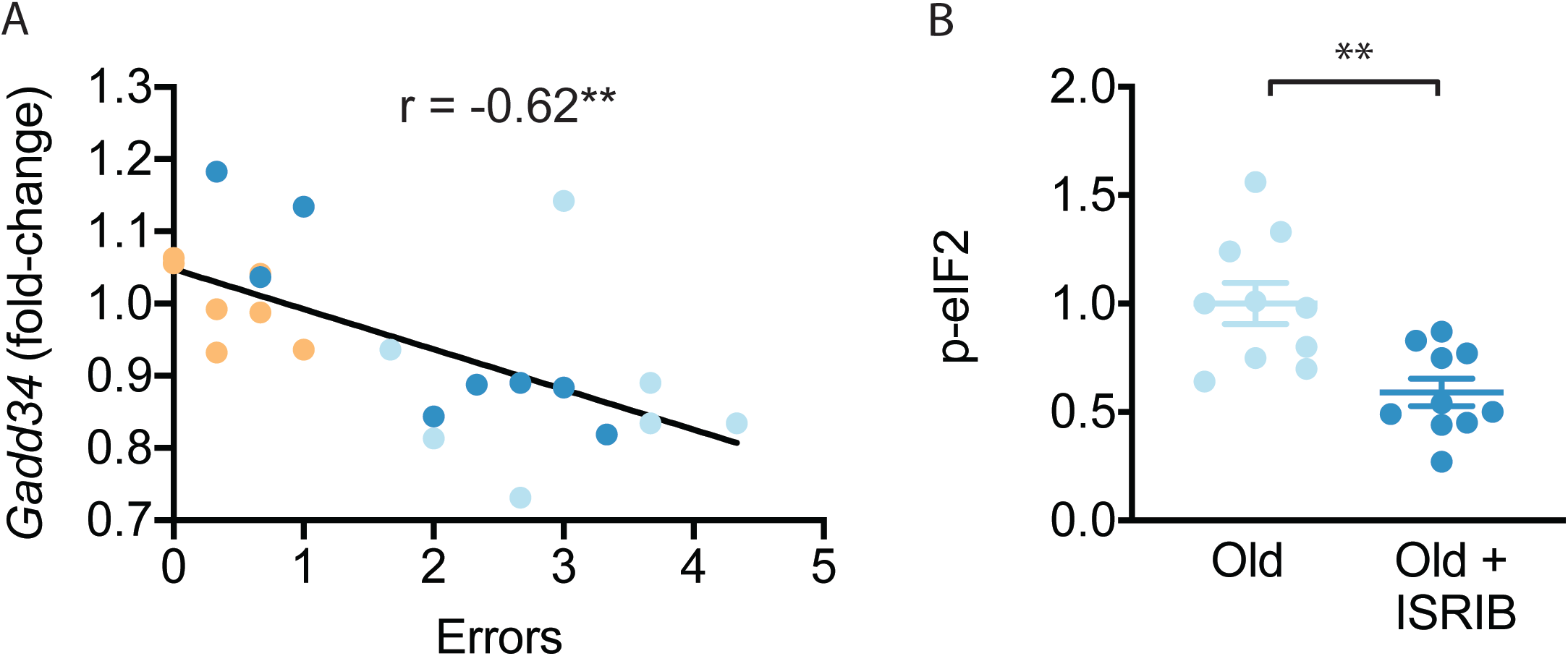
ISRIB treatment resets age-related ISR activation. (**A**) A significant negative correlation was measured between cognitive performance on day 2 of the RAWM (errors) and *Gadd34* mRNA expression. Linear regression was measured by Pearson R correlation, R value denoted with significance. (**B**) ISRIB treatment reduced p-eIF2α protein levels. Brain lysates of p-eIF2α protein levels normalized to actin. Old males n = 10; ISRIB males n = 10. Student’s t-test. **p < 0.01. Data are means ± SEM.

## CONCLUSION

We provide evidence for a direct involvement of the ISR in age-related cognitive decline. Temporary treatment with ISRIB causes down-regulation of ATF4 for at least 20 days. This “ISR reset” leads to improvement in spatial, working, and episodic memory. At a cellular level the cognitive enhancement is paralleled by i) improved intrinsic neuron excitability, ii) increased dendritic spine density, iii) reversal of age-induced changes in IFN and T cell responses in the hippocampus and blood, and iv) reversal of ISR activation. Thus, we identify broad-spectrum anatomical, cellular, and functional changes caused by ISR activation in old animals. If these findings in mice translate into human physiology, they offer hope and a tangible strategy to sustain cognitive ability as we age.

## Supporting information

All supplemental figures

---

**Supplemental Figure 1. ISRIB downregulates ATF4 during administration**. The impact of ISRIB on known ISR activation pathways was investigated by Western blot analysis of brain lysates after 3 ISRIB injections. (**A**) Raw western blot data. Each lane represents an individual animal brain extract. (**B**) ISRIB treatment reduced ATF4 protein levels during drug administration. Old males (7) and females (3): Old n = 10; Old + ISRIB n = 10. Student’s t-test. *p < 0.05. Individual animal values represented by dots; lines depict group mean ± SEM.

**Supplemental Figure 2. ISRIB down-regulates the ISR in the brain of old mice**. The impact of ISRIB on known ISR kinases and activation pathways was investigated by Western blot analysis of brain lysates (raw Western blot data) at day 20.(**A**) ATF4 (**B**) p-GCN2, p-PKR, p-PERK. Each lane represents an individual animal brain extract.

**Supplemental Figure 3. ISR inhibitors relieve age-induced deficits in spatial learning.** RAWM was used to measure age-induced deficits in spatial learning. Animals ran 2 blocks (3 trials/block) on each learning day. (**A**) Old animals performed significantly worse that young animals. Two-way repeated measures ANOVA revealed a significant interaction (p < 0.001). Bonferroni post-hoc to determine differences at various blocks. Old males n = 19, Young males n = 10. *p < 0.05. (**B**) ISRIB or vehicle administration (2.5 mg/kg intraperitoneal) occurred days 0-2. Compared with the old group, ISRIB treated animals made significantly fewer errors over the course of learning. Two-way repeated measures ANOVA reveals a significant difference between groups p < 0.05. Old males n = 19; Old + ISRIB males n = 15. (**C**) No differences were measured between young +/− ISRIB administration. Two-way repeated measures ANOVA revealed no significant differences. Young males n = 10; Young males + ISRIB n = 10 (**D**) Cmp-003 (5 mg/kg intraperitoneal) administration occurred days 0-2. Old mice that received Cmp-003 performed significantly better than old mice that received vehicle. Two-way repeated measures ANOVA revealed a significant group (p < 0.01) and time effect (p < 0.05). Old males n = 9, Old males + Cmp-003 n = 9. **p < 0.01. Data are means ± SEM.

**Supplemental Figure 4. ISRIB reduces age-induced memory deficits in female mice.** RAWM was used to measure age-induced deficits in learning and memory. ISRIB treatment improved memory one week after administration in female rodents. Student t-test. Old female n = 12; Old female + ISRIB n = 11. * p < 0.05. Individual animal scores represented by dots, lines depict group mean and SEM.

**Supplementary Figure 5. Age and ISRIB treatment do not modify other passive or active intrinsic membrane properties in CA1 pyramidal neurons.** (**A**) Representative traces from CA1 pyramidal neurons showing the membrane potential response to a 250 pA current injection in neurons from old animals treated with either vehicle (light blue) or ISRIB (dark blue) or young animals treated with vehicle (orange). Quantification of the action potential (AP) including the half width (**B**), amplitude (**C**), and threshold (**D**) did not show significant differences between CA1 pyramidal recordings from old, old + ISRIB-treated, or young mice. Likewise, evaluation of the maximum firing frequency (**E**) or how the frequency of spiking changes over time, quantified by the adaptation index (**F**) or with current injection, quantified by the slope of the relationship of firing frequency versus amplitude of current injection (F/I slope) (**G**) was also not significantly different between groups. Finally, passive membrane properties including the membrane time constant (tau) (**H**), membrane resistance (Rm) (**I**), and the resting membrane potential (**J**) were not significantly altered by age or ISRIB treatment. Each neuron is represented with a symbol; solid lines indicate the mean ± SEM. (One-way ANOVA for all comparisons; Neurons: and Young males n = 12 (5 animals); Old males n = 15 (5 animals), Old + ISRIB males n = 22 (7 animals) with 1-5 neurons recorded per animal.

**Supplemental Figure 6. Age and ISRIB treatment do not affect spontaneous excitatory post-synaptic currents (sEPSC) in CA1 pyramidal neurons.** (**A**) Representative whole cell voltage-clamp recordings showing sEPSCs from CA1 pyramidal neurons from old animals treated with either vehicle (light blue) or ISRIB (dark blue) or young animals treated with vehicle (orange). Arrows denote synaptic currents. (**B**) The sEPSC amplitude was not significantly difference between groups (one-way ANOVA). (**C**) The sEPSC frequency was unchanged after ISRIB treatment or compared to young mice (Kruskal-Wallis test). The median amplitude or frequency for each neuron is represented with a symbol; solid lines indicate the mean ± SEM. (Neurons: Young males n = 11 (5 animals); Old males n = 15 (5 animals), Old + ISRIB males n = 18 (7 animals) with 1-5 neurons recorded per animal.)

**Supplemental Figure 7. ISRIB administration does not impact glial cell activation.** (**A**) ISRIB administration scheme. (**B-G**) Glial cell was quantified in the stratum radiatum of the CA1 region of the dorsal hippocampus from old Thy1-YFP-H mice. GFAP was used to measure astrocyte activity. Representative images for GFAP staining of (**B**) old and (**C**) old + ISRIB mouse. (**D**) No differences in GFAP percent area were measured when comparing old and old + ISRIB animals. Iba-1 was used to measure microglia activity. Representative images for Iba-1 staining of (**E**) old and (**F**) old + ISRIB mouse. (**G**) No differences in Iba-1 percent area were measured when comparing old and old + ISRIB animals. 63x magnification with a water immersion objective. Old males n = 12 - 19 slides (3 mice); Old males + ISRIB n = 18 - 20 slides (4 mice). Individual slide scores (relative to old mice) represented in dots, lines depict group mean ± SEM.

**Supplemental Figure 8. ISRIB treatment breaks age-related ISR activation**. The impact of ISRIB on p-eIF2 was investigated by Western blot analysis of brain lysates after 3 ISRIB injections. Raw western blot data. Each lane represents an individual animal brain extract.

## METHODS

### Animals

All experiments were conducted in accordance with National Institutes of Health (NIH) Guide for the Care and Use of Laboratory Animals and approved by the Institutional Animal Care and Use Committee of the University of California, San Francisco (Protocol 170302). Male and female C57B6/J wild-type (WT) mice were received from the National Institute of Aging. Thy-1-YFP-H (in C57 background) were bred and aged in house. Old animals started experimentation at ~19 months of age and young animals 3-6 months of age. Animal shipments were received at least one week prior to start of experimentation to allow animals to habituation the new surroundings. Mice were group housed in environmentally controlled conditions with reverse light cycle (12:12 h light:dark cycle at 21 ± 1 °C; ~50% humidity) and provided food and water ad libitum. Behavioral analysis was performed during the dark cycle.

### Drug Administration

ISRIB solution was made by dissolving 5 mg ISRIB in 2.5 mLs dimethyl sulfoxide (DMSO) (PanReac AppliChem, 191954.1611). The solution was gently heated in a 40 °C water bath and vortexed every 30 s until the solution became clear. Next 1 mL of Tween 80 (Sigma Aldrich, P8074) was added, the solution was gently heated in a 40 °C water bath and vortexed every 30 s until the solution became clear. Next, 10 mL of polyethylene glycol 400 (PEG400) (PanReac AppliChem, 142436.1611) solution was added gently heated in a 40 °C water bath and vortexed every 30 s until the solution became clear. Finally, 36.5 mL of 5% dextrose (Hospira, RL-3040) was added. The solution was kept at room temperature throughout the experiment. Each solution was used for injections up to 7 day maximum. The vehicle solution consisted of the same chemical composition and concentration (DMSO, Tween 80, PEG400 and 5% dextrose). Stock ISRIB solution was at 0.1 mg/ml and injections were at 2.5 mg/kg. Each animal received an intraperitoneal injection of 2.5x their body weight.

Cmp-003 solution was made by dissolving Cmp-003 (donated by Praxis Biotech) in 50% PEG400 (PanReac AppliChem, 142436.1611) and 50% sterile water. The solution was gently heated in a 40 °C water bath and vortexed every 30 s until the solution became clear. Stock Cmp-003 solution was at 0.5 mg/ml and animal injections were at 5.0 mg/kg. Solution was used immediately and made fresh daily.

### Behavioral assessment of cognitive functions

For all behavioral assays the experimenter(s) were blinded to therapeutic intervention. Prior to behavioral analysis animals were inspected for gross motor impairments. Animals were inspected for whisker loss, limb immobility (included grip strength) and eye occlusions. If animals displayed *any* of these impairments, they were excluded. Behavioral assessment was recorded and scored using a video tracking and analysis setup (Ethovision XT 8.5, Noldus Information Technology).

#### Radial Arm Water Maze

The radial arm water maze (RAWM) was used to test spatial learning and memory in rodents (*27*, *42*). The pool is 118.5 cm in diameter with 8 arms, each 41 cm in length, and an escape platform. The escape platform is slightly submerged below the water level, so it is not visible to the animals. The pool was filled with water that was rendered opaque by adding white paint (Crayola, 54–2128-053). Visual cues are placed around the room such that they were visible to animals exploring the maze. Animals ran 6 trials a day during learning and 3 trials during each memory probe. On both learning and memory days there is a 10-minute inter-trial interval. Animals were trained for 2 days and then tested on memory tests 24 hours and 8 days after training. During a trial, animals were placed in a random arm that did not include the escape platform. Animals were allowed 1 min to locate the escape platform. On successfully finding the platform, animals remained there for 10 s before being returned to their warmed, holding cage. On a failed trial, animals were guided to the escape platform and then returned to their holding cage 10 s later. The escape platform location was the same, whereas the start arm varied between trials.

Animals were injected (intraperitoneal) with either vehicle or ISRIB (2.5 mg/kg) starting the day prior to behavior (**Figure 2A**) and after each of the final trials of the learning days (day 1 and 2) for a total of three doses. No injections were given when memory was tested on days 3 and 10. RAWM data were collected through a video tracking and analysis setup (Ethovision XT 8.5, Noldus Information Technology). The program automatically analyzed the number of entries into non-target arms made per trial. Every three trials were averaged into a block to account for large variability in performance; each learning day thus consisted of 2 blocks, whereas each memory test was one block each. Importantly, in all animal cohorts tested (regardless of age or drug treatment) learning was measured (Significant time effect observed in all Two-way repeated measure ANOVA analysis when groups are analyzed independently).

#### Delayed Matching to Place Barnes Maze

Beginning at day 20 animals were tested on DMP using a modified Barnes maze (*27*, *44*). The maze consisted of a round table 112 cm in diameter with 40 escape holes arranged in three concentric rings consisting of 8, 16, and 16 holes at 20, 35, and 50 cm from the center of the maze, respectively. An escape tunnel was connected to one of the outer holes. Visual cues were placed around the room such that they were visible to animals on the table. Bright overhead lighting and a loud tone (2 KHz, 85 db) were used as aversive stimuli to motivate animals to locate the escape tunnel. The assay was performed for 4 days (days 20-23). The escape tunnel location was moved for each day and animals ran four trials on the first two days and 3 trials on the last two days. During a trial, animals were placed onto the center of the table covered by an opaque plastic box so they were not exposed to the environment. After they had been placed on the table for 10 s, the plastic box was removed and the tone started playing, marking the start of the trial. Animals were given 90 s to explore the maze and locate the escape tunnel. When the animals successfully located and entered the escape tunnel, the tone was stopped. If the animals failed to find the escape tunnel after 90 s, they were guided to the escape tunnel before the tone was stopped. Animals remained in the escape tunnel for 10 s before being returned to their home cage. The maze was cleaned with ethanol between each trial. A new escape tunnel was used for each trial. The experimenter was blind to the treatment groups during the behavioral assay. Each trial was recorded using a video tracking and analysis setup (Ethovision XT 8.5, Noldus Information Technology) and the program automatically analyzed the amount of time required to locate the escape tunnel. Animal improvement was calculated by Day 20 escape latency – Day 23 escape latency.

### Tissue collection

All mice were lethally overdosed using a mixture of ketamine (10 mg/ml) and xylaxine (1 mg/ml). Once animals were completely anesthetized, blood was extracted by cardiac puncture and animals were perfused with 1X phosphate buffer solution, pH 7.4 (Gibco, Big Cabin, OK, -70011-044) until the livers were clear (~1–2 min). For Western blot analysis following phosphate buffered solution (PBS), the whole brain (regions dissected discussed below) was rapidly removed and snap frozen on dry ice and stored at −80 °C until processing.

### Western Blot Analysis

Animals received all 3 ISRIB injections and were terminated 20 h after the third injection (as described above). Frozen brain lysates or hippocampi isolates were then homogenized with a T 10 basic ULTRA-TURRAX (IKa) in ice-cold buffer lysis (Cell Signaling 9803) and protease and phosphatase inhibitors (Roche). Lysates were sonicated for 3 min and centrifuged at 13,000 rpm for 20 min at 4°C. Protein concentration in supernatants was determined using BCA Protein Assay Kit (Pierce). Equal amount of proteins was loaded on SDS-PAGE gels. Proteins were transferred onto 0.2 μm PVDF membranes (BioRad) and probed with primary antibodies diluted in Tris-buffered saline supplemented with 0.1% Tween 20 and 3% bovine serum albumin.

ATF4 (11815) (Cell Signaling), p-GCN2 (Abcam Cat No ab-75836), p-PERK (Cell Signaling Cat No 3179), p-PKR (Abcam Cat No ab-32036), and p-eIF2 (Cell Signaling, Cat No 3597) and β-actin (Sigma-Aldrich) antibodies were used as primary antibodies. HRP-conjugated secondary antibodies (Rockland) were employed to detect immune-reactive bands using enhanced chemiluminescence (ECL Western Blotting Substrate, Pierce) according to the manufacturer instructions. Quantification of protein bands was done by densitometry using ImageJ software.

ATF4, p-GCN2, p-PERK, p-PKR and p-eIF2 levels were normalized to β-actin expression and fold-change was calculated as the levels relative to the expression in vehicle-treated derived samples, which corresponds to 1.

### Electrophysiology

Sagittal brain slices (250 μm) including the hippocampus were prepared from old mice (~19 mo) treated with either vehicle or ISRIB or young mice (~3 mo), treated with vehicle, 12-18 hours prior (n = 5, 7, and 5 per group respectively). Mice were anesthetized with Euthasol (0.1 ml / 25 g, Virbac, Fort Worth, TX, NDC-051311-050-01), and transcardially perfused with an ice-cold sucrose cutting solution containing (in mM): 210 sucrose, 1.25 NaH_2_PO_4_, 25 NaHCO_3_, 2.5 KCl, 0.5 CaCl_2_, 7 MgCl_2_, 7 dextrose, 1.3 ascorbic acid, 3 sodium pyruvate (bubbled with 95% O_2_ − 5% CO_2_, pH 7.4) (see **Supplemental Table 1** for reagent information). Mice were then decapitated and the brain was isolated in the same sucrose solution and cut on a slicing vibratome (Leica, VT1200S, Leica Microsystems, Wetzlar, Germany). Slices were incubated in a holding solution (composed of (in mM): 125 NaCl, 2.5 KCl, 1.25 NaH_2_PO_4_, 25 NaHCO_3_, 2 CaCl_2_, 2 MgCl_2_, 10 dextrose, 1.3 ascorbic acid, 3 sodium pyruvate, bubbled with 95% O_2_ − 5% CO_2_, pH 7.4) at 36 °C for 30 min and then at room temperature for at least 30 min until recording.

Whole cell recordings were obtained from these slices in a submersion chamber with a heated (32 – 34 °C) artificial cerebrospinal fluid (aCSF) containing (in mM): 125 NaCl, 3 KCl, 1.25 NaH_2_PO_4_, 25 NaHCO_3_, 2 CaCl_2_, 1 MgCl_2_, 10 dextrose (bubbled with 95% O_2_ - 5% CO_2_, pH 7.4). Patch pipettes (3–6 MΩ) were manufactured from filamented borosilicate glass capillaries (Sutter Instruments, Novato, CA, BF100-58-10) and filled with an intracellular solution containing (in mM): 135 KGluconate, 5 KCl, 10 HEPES, 4 NaCl, 4 MgATP, 0.3 Na_3_GTP, 7 2K-phosphcreatine, and 1-2% biocytin. CA1 pyramidal neurons were identified using infrared microscopy with a 40x water-immersion objective (Olympus, Burlingame, CA). Recordings were made using a Multiclamp 700B (Molecular Devices, San Jose, CA) amplifier, which was connected to the computer with a Digidata 1440A ADC (Molecular Devices, San Jose, CA), and recorded at a sampling rate of 20 kHz with pClamp software (Molecular Devices, San Jose, CA). We did not correct for the junction potential, but access resistance and pipette capacitance were appropriately compensated before each recording.

The passive membrane and active action potential spiking characteristics were assessed by injection of a series of hyperpolarizing and depolarizing current steps with a duration of 250 ms from −250 pA to 700 nA (in increments of 50 pA). The resting membrane potential was the measured voltage of the cell 5 min after obtaining whole cell configuration without current injection. A holding current was then applied to maintain the neuron at −67 +/− 2 mV before/after current injections. The input resistance was determined from the steady-state voltage reached during the −50 pA current injection. The membrane time constant was the time required to reach 63% of the maximum change in voltage for the −50 pA current injection. Action potential parameters including the half width, threshold, and amplitude were quantified from the first action potential elicited. Action potential times were detected by recording the time at which the positive slope of the membrane potential crossed 0 mV. From the action potential times, the instantaneous frequency for each action potential was determined (1 / inter spike interval). The maximum firing frequency was the highest frequency of firing identified throughout all current injections. Action potential rate as a function of current injection was examined by plotting the first instantaneous action potential frequency versus current injection amplitude. The F/I slope was then determined from the best linear fit of the positive values of this plot. The action potential or spike threshold was defined as the voltage at which the third derivative of V (d3V/dt) was maximal just prior to the action potential peak. The action potential (AP) amplitude was calculated by measuring the voltage difference between the peak voltage of the action potential and the spike threshold. The half-width of the action potential was determined as the duration of the action potential at half the amplitude. The adaptation index of each cell was the ratio of the last over the first instantaneous firing frequency, calculated at 250 pA above the current step that first elicited spiking. The afterhyperpolarization (AHP) was calculated as the change in voltage from baseline (measured as the mean voltage over a 100 ms interval 600 ms after termination of a current injection that first elicited at least 12 spikes corresponding to a firing frequency of ~50 Hz) compared to immediately after cessation of current injection (the minimum voltage reached in the first 175 ms immediately after cessation of current injection). Cells were excluded from analysis if excessive synaptic input was noted during recording of the mAHP or if the cell did not fire at least 12 spikes during current injections.

To measure the spontaneous excitatory postsynaptic currents (sEPSCs), cells were recorded in voltage clamp at a holding potential of −75 mV for 4 min, a holding potential that should have little inhibitory components given the reversal potential of chloride with these solutions. Analysis of sEPSCs was performed using a template matching algorithm in ClampFit 10.7 (Molecular Devices, San Jose, CA). The template was created using recordings from multiple pyramidal cells and included several hundred synaptic events. Access resistance (Ra) was monitored during recordings, and recordings were terminated if Ra exceeded 30 megaohms. Only stable recordings (< 50 pA baseline change) with a low baseline noise (< 8 pA root mean square) were included. The first 250 synaptic events or all the events measured in the 4 min interval from each cell were included for analysis.

### Fluorescent spine imaging preparation

For fluorescent spine analysis, following PBS animals were perfused with ice-cold 4% paraformaldehyde, pH 7.5 (PFA, Sigma Aldrich, St. Louis, MO, 441244) and fixed for 4 - 24 h followed by sucrose (Fisher Science Education, Nazareth, PA, S25590A) protection (15% to 30%). Brains were embedded with 30% sucrose/ Optimal Cutting Temperature Compound (Tissue Tek, Radnor, PA, 4583) mixture on dry ice and stored at −80 °C. Brains were sectioned into 20 μm slides using a Leica cryostat (Leica Microsystems, Wetzlar, Germany) and mounted on slides (ThermoFisher Scientific, South San Francisco, CA). Slides were brought to room temperature (20 °C) prior to use. Tissues were fixed using ProLong Gold (Invitrogen, Carlsbad, CA, P36930) and a standard slide cover sealed with nail polish.

### Spine density quantification

For spine density quantification, whole brains from young and old male Thy1-YFP-H transgenic line were used. 3-6 images separated by 60-140 μm in the dorsal hippocampus were imaged per animal and used for dendritic spine density analysis. 9.3 μm z-stack images were acquired on a Zeiss Laser-Scanning Confocal microscope (Zeiss LSM 780 NLO FLIM) at the HDFCCC Laboratory for Cell Analysis Shared Resource Facility. 63x magnification with a water immersion objective. All protrusions from the dendrites were manually counted as spines regardless of morphology. Two individuals (blinded to age and treatment) analyzed a total length of at least 3200 μm of dendrites from each animal using NIH FIJI analysis software (v1.52n). Individual dendritic spine was calculated as density per micron and graphed relative to old mice.

### qPCR Analysis

Hippocampus samples, of approximately the same size per animal were process as previously described (*50*, *51*). Relative gene expression was determined using the 2^−ΔΔCt^ method and normalized using GAPDH. Primers used were the following:

*Cd3*: Fw 5’ TGACCTCATCGCAACTCTGCTC-3’ Rev 5’ TCAGCAGTGCTTGAACCTCAGC-3’
*Ifit1*: Fw 5’ CTGAGATGTCACTTCACATGGAA-3’ Rev 5’ GTGCATCCCCAATGGGTTCT-3’
*Rtp4*: Fw 5’ TGGGAGCAGACATTTCAAGAAC-3’, Rev 5’ACCTGAGCAGAGGTCCAACTT-3’
*Gbp10*: Fw 5’ GGAGGCTCAAGAGAAAAGTCACA-3’, Rev 5’ AAGGAAAGCCTTTTGATCCTTCAGC-3’
*Ccl2*: Fw 5’ GCTGACCCCAAGAAGGAATG-3’ Rev 5’ GTGCTTGAGGTGGTTGTGGA-3’
*Il1β*: Fw 5’ TGTAATGAAAGACGGCACACC-3’ Rev 5’ TCTTCTTTGGGTATTGCTTGG-3’
*Tnfα*: Fw 5’ TGCCTATGTCTCAGCCTCTTC-3’ Rev 5’ GAGGCCATTTGGGAACTTCT-3’
*Il-6*: Fw 5’ TACCACTTCACAAGTCGGAGGC-3’ Rev 5’ CTGCAAGTGCATCATCGTTGTTC-3’
*Irf7:* Fw 5’- GAGACTGGCTATTGGGGGAG-3’ Rev 5’- GACCGAAATGCTTCCAGGG-3’
*Ifitm3:* Fw 5’- CCCCCAAACTACGAAAGAATCA-3’ Rev 5’- ACCATCTTCCGATCCCTAGAC-3’
*Isg15:* Fw 5’- GGTGTCCGTGACTAACTCCAT-3’ Rev 5’- TGGAAAGGGTAAGACCGTCCT-3’
*Ifi204:* Fw 5’- AGCTGATTCTGGATTGGGCA-3’ Rev 5’- GTGATGTTTCTCCTGTTACTTCTGA-3’
*Eif2ak2 (Pkr):* Fw 5’- CTGGTTCAGGTGTCACCAAAC-3’ Rev 5’- ACAACGCTAGAGGATGTTCCG-3’
*Cd11b*: Fw 5’- CTGAGACTGGAGGCAACCAT- 3’ Rev 5’ GATATCTCCTTCGCGCAGAC-3’
*Il-10*: Fw 5’- GCCAAGCCTTATCGGAAATG- 3’ Rev 5’ CACCCAGGGAATTCAAATGC-3’
*Bdnf*: Fw 5’- GGCTGACACTTTTGAGCACGT - 3’ Rev 5’ CTCCAAAGGCACTTGACTGCTG -3’
*Ophn1*: Fw 5’- CTTCCAGGACAGCCAACCATTG- 3’ Rev 5’ CTTAGCACCTGGCTTCTGTTCC -3’
*Gbs5*: Fw 5’- CTGAACTCAGATTTTGTGCAGGA - 3’ Rev 5’ CATCGACATAAGTCAGCACCAG -3’
*Oasl1*: Fw 5’- CAGGAGCTGTACGGCTTCC - 3’ Rev 5’ CCTACCTTGAGTACCTTGAGCAC -3’
*Gadd34*: Fw 5’- GGCGGCTCAGATTGTTCAAAGC - 3’ Rev 5’ CCAGACAGCAAGGAAATGGACTG -3’
*Gapdh*: Fw 5’ AAATGGTGAAGGTCGGTGTG-3’ Rev 5’ TGAAGGGGTCGTTGATGG-3’

### Flow Cytometric Analysis

To assess circulating cell populations peripheral blood was collected by cardiac puncture and transferred into an EDTA collection tube. Blood was aliquoted into flow cytometry staining tubes and stained with surface antibodies for 30-60 min at room temperature (*52*). Surface antibodies included anti-CD45 (FITC-conjugated; BD Biosciences), Ly-6G (PE-conjugated; BD Biosciences), CD8 (PE-Cy7-conjugated; BD Biosciences), CD4 (APC-conjugated; BD B), and CD11b (APC-Cy7; BD Biosciences). Leukocyte subpopulations were identified as follows: Forward and side scatter was used to exclude debris and doublet populations. Specific T-cell populations were identified as follows: CD4 T-cell subsets were CD4+, CD45+, Ly-6G−, CD8−, CD11b−. CD8 T-cell subsets were CD8+, CD45+, Ly-6G−, CD4−,CD11b−. After surface antibody staining, red blood cells were lysed with RBC lysis (BD Biosciences). Cell population gating occurred as previously described (*52*). (Data were collected on an LSRII (BD) and analyzed with Flowjo™ software (v10, Tree Star Inc.).

### Statistical Analysis

Results were analyzed using Prism software or IBM SPSS Statistics. Individual animals. Individual animal scores represented by dots, lines depict group mean and SEM. Student t-test, one-way ANOVA, two-way repeated measures ANOVA and Pearson R correlations were used (individual statistical tool and post-hoc analysis denoted in Figure Legends). p values of < 0.05 considered as significant.

**Supplemental Table 1.**
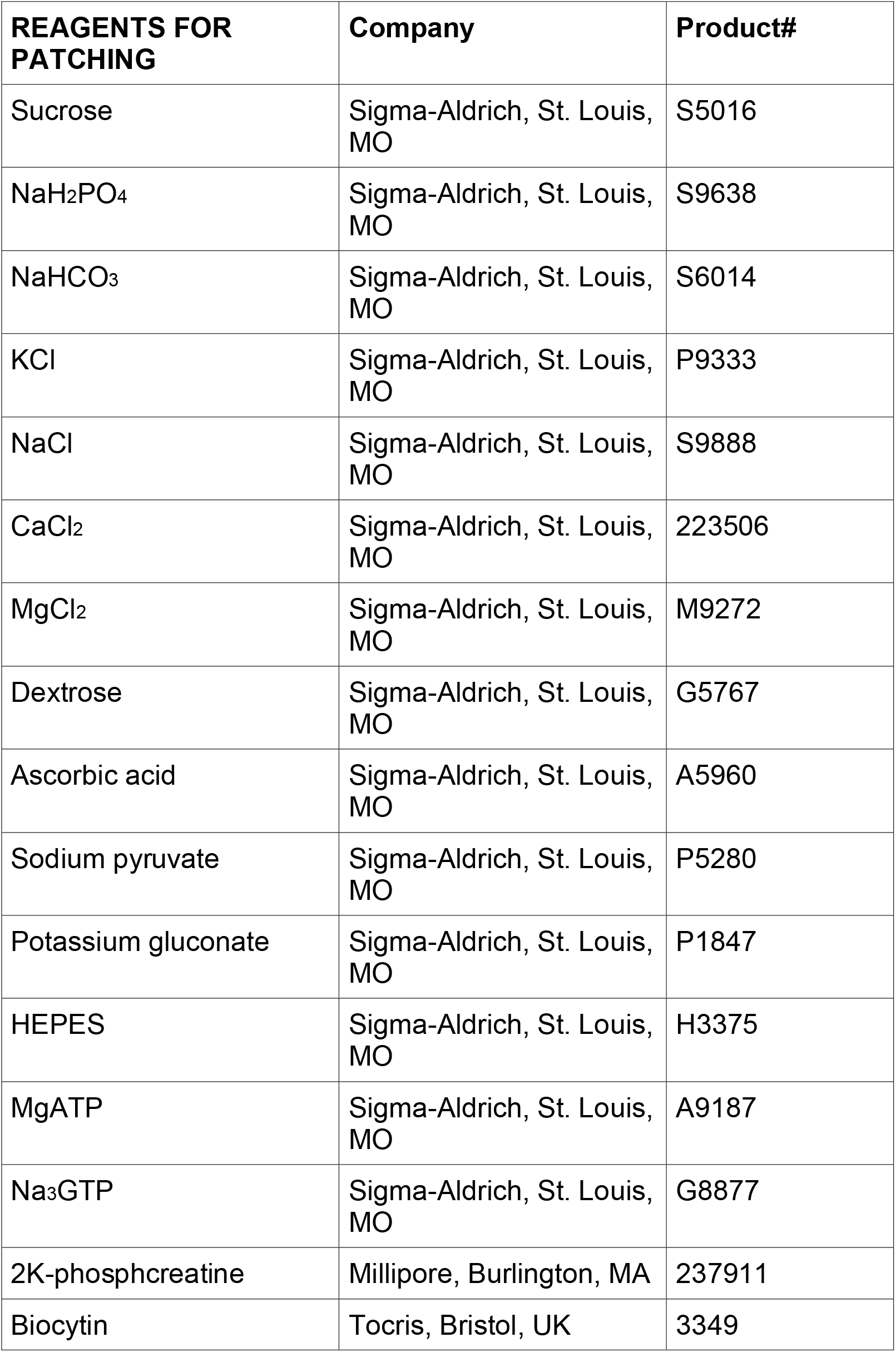
List of Electrophysiology Reagents.

| **REAGENTS FOR PATCHING** | **Company** | **Product#** |
| --- | --- | --- |
| Sucrose | Sigma-Aldrich, St. Louis, MO | S5016 |
| NaH<sub>2</sub>PO<sub>4</sub> | Sigma-Aldrich, St. Louis, MO | S9638 |
| NaHCO<sub>3</sub> | Sigma-Aldrich, St. Louis, MO | S6014 |
| KCl | Sigma-Aldrich, St. Louis, MO | P9333 |
| NaCl | Sigma-Aldrich, St. Louis, MO | S9888 |
| CaCl<sub>2</sub> | Sigma-Aldrich, St. Louis, MO | 223506 |
| MgCl<sub>2</sub> | Sigma-Aldrich, St. Louis, MO | M9272 |
| Dextrose | Sigma-Aldrich, St. Louis, MO | G5767 |
| Ascorbic acid | Sigma-Aldrich, St. Louis, MO | A5960 |
| Sodium pyruvate | Sigma-Aldrich, St. Louis, MO | P5280 |
| Potassium gluconate | Sigma-Aldrich, St. Louis, MO | P1847 |
| HEPES | Sigma-Aldrich, St. Louis, MO | H3375 |
| MgATP | Sigma-Aldrich, St. Louis, MO | A9187 |
| Na<sub>3</sub>GTP | Sigma-Aldrich, St. Louis, MO | G8877 |
| 2K-phosphocreatine | Millipore, Burlington, MA | 237911 |
| Biocytin | Tocris, Bristol, UK | 3349 |

## ACKNOWLEDGEMENTS

This work was supported by the generous support of the Rogers Family (to S.R. and P.W.), the UCSF Weill Innovation Award (to S.R. and P.W.), the NIH/National Institute on Aging Grant R01AG056770 (to S.R.), the NRSA post-doctoral fellowship from the NIA F32AG054126 (to K.K), the National Institute for General Medicine (NIGMS) Initiative for Maximizing Student Development (R25GM056847) and the National Science Foundation (NSF) Graduate Fellowship Program (To E.S.F), the UCSF Clinical and National Center for Advanced Translational Sciences at NIH (UCSF-CTSI Grant Number TL1 TR001871) and the NIH/NINDS (K08NS114170) (To A.N), the Programa de Apoyo a Centros con Financiamiento Basal AFB 170004 (to S.B.). P.W. is an Investigator of the Howard Hughes Medical Institute.

We thank Dr. Vikaas Sohal for providing equipment for electrophysiological recordings and advice on analysis. We thank Dr. Spyros Darmanis and Rene Sit from the Chan Zuckerberg Biohub for their assistance with analysis.

The authors would like to thank Praxis Biotech LLC, San Francisco, CA for providing samples of Cmp-003, for use in experiments described in this publication.

Microscopic imaging was obtained at the HDFCCC Laboratory for Cell Analysis Shared Resource Facility which is funded through grants from NIH (P30CA082103 and S10 ODo21818-01).

## REFERENCES

1. Cognitive Aging. Progress in Understanding and Opportunities for Action (2015).

2. S. L. Connelly, L. Hasher, R. T. Zacks, Age and reading: the impact of distraction. Psychol Aging 6, 533–541 (1991).

3. N. D. Anderson, F. I. Craik, M. Naveh-Benjamin, The attentional demands of encoding and retrieval in younger and older adults: 1. Evidence from divided attention costs. Psychol Aging 13, 405–423 (1998).

4. A. F. Kramer, S. Hahn, D. Gopher, Task coordination and aging: explorations of executive control processes in the task switching paradigm. Acta Psychol (Amst) 101, 339–378 (1999).

5. N. J. Cepeda, A. F. Kramer, J. C. Gonzalez de Sather, Changes in executive control across the life span: examination of task-switching performance. Dev Psychol 37, 715–730 (2001).

6. An Aging Nation: The Older Population in the United States (2014).

7. A. Chou, K. Krukowski, J. M. Morganti, L. K. Riparip, S. Rosi, Persistent Infiltration and Impaired Response of Peripherally-Derived Monocytes after Traumatic Brain Injury in the Aged Brain. Int J Mol Sci 19, (2018).

8. H. Yousef et al., Aged blood impairs hippocampal neural precursor activity and activates microglia via brain endothelial cell VCAM1. Nat Med 25, 988–1000 (2019).

9. S. A. Villeda et al., The ageing systemic milieu negatively regulates neurogenesis and cognitive function. Nature 477, 90–94 (2011).

10. J. M. Castellano et al., Human umbilical cord plasma proteins revitalize hippocampal function in aged mice. Nature 544, 488–492 (2017).

11. S. A. Villeda et al., Young blood reverses age-related impairments in cognitive function and synaptic plasticity in mice. Nat Med 20, 659–663 (2014).

12. F. T. Cabral-Miranda, G. Martinez, G. Medinas, D. Gerakis, Y. Miedema, T. Duran-Aniotz, C. Ardiles, AO. Gonzalez, C. Sabusap, C. Bermedo-Garcia, F. Adamson, S. Vitangcol, K. Huerta, H. Zhang, X. Nakamura, T. Pablo Sardi, S. Lipton, SA. Kenedy, BK. Cárdenas, JC. Palacios, AG. Plate, L. Henriquez, JP. Hetz, C., Control of mammalian brain aging by the unfolded protein response (UPR). Biorxiv, (2020).

13. J. F. Disterhoft, M. M. Oh, Alterations in intrinsic neuronal excitability during normal aging. Aging Cell 6, 327–336 (2007).

14. E. C. McKiernan, D. F. Marrone, CA1 pyramidal cells have diverse biophysical properties, affected by development, experience, and aging. PeerJ 5, e3836 (2017).

15. M. M. Oh, F. A. Oliveira, J. F. Disterhoft, Learning and aging related changes in intrinsic neuronal excitability. Front Aging Neurosci 2, 2 (2010).

16. V. Rizzo, J. Richman, S. V. Puthanveettil, Dissecting mechanisms of brain aging by studying the intrinsic excitability of neurons. Front Aging Neurosci 6, 337 (2014).

17. L. A. Schimanski, C. A. Barnes, Neural Protein Synthesis during Aging: Effects on Plasticity and Memory. Front Aging Neurosci 2, (2010).

18. V. Azzu, T. G. Valencak, Energy Metabolism and Ageing in the Mouse: A Mini-Review. Gerontology 63, 327–336 (2017).

19. C. Franceschi et al., Inflamm-aging. An evolutionary perspective on immunosenescence. Ann N Y Acad Sci 908, 244–254 (2000).

20. K. Baruch et al., Aging. Aging-induced type I interferon response at the choroid plexus negatively affects brain function. Science 346, 89–93 (2014).

21. B. W. Dulken et al., Single-cell analysis reveals T cell infiltration in old neurogenic niches. Nature 571, 205–210 (2019).

22. J. B. Flexner, L. B. Flexner, E. Stellar, G. De La Haba, R. B. Roberts, Inhibition of protein synthesis in brain and learning and memory following puromycin. J Neurochem 9, 595–605 (1962).

23. C. Lopez-Otin, M. A. Blasco, L. Partridge, M. Serrano, G. Kroemer, The hallmarks of aging. Cell 153, 1194–1217 (2013).

24. M. C. Ingvar, P. Maeder, L. Sokoloff, C. B. Smith, Effects of ageing on local rates of cerebral protein synthesis in Sprague-Dawley rats. Brain 108 (Pt 1), 155–170 (1985).

25. C. B. Smith, Y. Sun, L. Sokoloff, Effects of aging on regional rates of cerebral protein synthesis in the Sprague-Dawley rat: examination of the influence of recycling of amino acids derived from protein degradation into the precursor pool. Neurochem Int 27, 407–416 (1995).

26. H. P. Harding et al., An integrated stress response regulates amino acid metabolism and resistance to oxidative stress. Mol Cell 11, 619–633 (2003).

27. A. Chou et al., Inhibition of the integrated stress response reverses cognitive deficits after traumatic brain injury. Proc Natl Acad Sci U S A 114, E6420–E6426 (2017).

28. K. Krukowski et al., Integrated Stress Response Inhibitor Reverses Sex-Dependent Behavioral and Cell-Specific Deficits after Mild Repetitive Head Trauma. J Neurotrauma, (2020).

29. M. Costa-Mattioli, P. Walter, The integrated stress response: From mechanism to disease. Science 368, (2020).

30. C. C. Kaczorowski, J. F. Disterhoft, Memory deficits are associated with impaired ability to modulate neuronal excitability in middle-aged mice. Learn Mem 16, 362–366 (2009).

31. O. von Bohlen und Halbach, C. Zacher, P. Gass, K. Unsicker, Age-related alterations in hippocampal spines and deficiencies in spatial memory in mice. J Neurosci Res 83, 525–531 (2006).

32. B. Xu et al., Loss of thin spines and small synapses contributes to defective hippocampal function in aged mice. Neurobiol Aging 71, 91–104 (2018).

33. E. B. Bloss et al., Evidence for reduced experience-dependent dendritic spine plasticity in the aging prefrontal cortex. J Neurosci 31, 7831–7839 (2011).

34. N. Yasumatsu, M. Matsuzaki, T. Miyazaki, J. Noguchi, H. Kasai, Principles of long-term dynamics of dendritic spines. J Neurosci 28, 13592–13608 (2008).

35. U. I. Onat et al., Intercepting the Lipid-Induced Integrated Stress Response Reduces Atherosclerosis. J Am Coll Cardiol 73, 1149–1169 (2019).

36. A. Deczkowska et al., Mef2C restrains microglial inflammatory response and is lost in brain ageing in an IFN-I-dependent manner. Nat Commun 8, 717 (2017).

37. A. G. Hinnebusch, I. P. Ivanov, N. Sonenberg, Translational control by 5'-untranslated regions of eukaryotic mRNAs. Science 352, 1413–1416 (2016).

38. N. Sonenberg, A. G. Hinnebusch, Regulation of translation initiation in eukaryotes: mechanisms and biological targets. Cell 136, 731–745 (2009).

39. A. Chen et al., Inducible enhancement of memory storage and synaptic plasticity in transgenic mice expressing an inhibitor of ATF4 (CREB-2) and C/EBP proteins. Neuron 39, 655–669 (2003).

40. S. Pasini, C. Corona, J. Liu, L. A. Greene, M. L. Shelanski, Specific downregulation of hippocampal ATF4 reveals a necessary role in synaptic plasticity and memory. Cell Rep 11, 183–191 (2015).

41. R. C. Wek, Role of eIF2alpha Kinases in Translational Control and Adaptation to Cellular Stress. Cold Spring Harb Perspect Biol 10, (2018).

42. J. Alamed, D. M. Wilcock, D. M. Diamond, M. N. Gordon, D. Morgan, Two-day radial-arm water maze learning and memory task; robust resolution of amyloid-related memory deficits in transgenic mice. Nat Protoc 1, 1671–1679 (2006).

43. A. M. Horowitz et al., Blood factors transfer beneficial effects of exercise on neurogenesis and cognition to the aged brain. Science 369, 167–173 (2020).

44. X. Feng, K. Krukowski, T. Jopson, S. Rosi, Delayed-matching-to-place Task in a Dry Maze to Measure Spatial Working Memory in Mice. Bio Protoc 7, (2017).

45. M. A. Oliveira Pisco A, Schaum N, Karkanias J, Neff NF, Darmanis S, Wyss-Coray T, Quake SR, A Single Cell Transcriptomic Atlas Characterizes Aging Tissues in the Mouse. bioRxiv, (2020).

46. D. Mrdjen et al., High-Dimensional Single-Cell Mapping of Central Nervous System Immune Cells Reveals Distinct Myeloid Subsets in Health, Aging, and Disease. Immunity 48, 599 (2018).

47. M. H. Brush, D. C. Weiser, S. Shenolikar, Growth arrest and DNA damage-inducible protein GADD34 targets protein phosphatase 1 alpha to the endoplasmic reticulum and promotes dephosphorylation of the alpha subunit of eukaryotic translation initiation factor 2. Mol Cell Biol 23, 1292–1303 (2003).

48. J. H. Connor, D. C. Weiser, S. Li, J. M. Hallenbeck, S. Shenolikar, Growth arrest and DNA damage-inducible protein GADD34 assembles a novel signaling complex containing protein phosphatase 1 and inhibitor 1. Mol Cell Biol 21, 6841–6850 (2001).

49. I. Novoa, H. Zeng, H. P. Harding, D. Ron, Feedback inhibition of the unfolded protein response by GADD34-mediated dephosphorylation of eIF2alpha. J Cell Biol 153, 1011–1022 (2001).

50. K. Krukowski et al., Traumatic Brain Injury in Aged Mice Induces Chronic Microglia Activation, Synapse Loss, and Complement-Dependent Memory Deficits. Int J Mol Sci 19, (2018).

51. K. Krukowski et al., Female mice are protected from space radiation-induced maladaptive responses. Brain Behav Immun 74, 106–120 (2018).

52. K. Krukowski, T. Jones, M. Campbell-Beachler, G. Nelson, S. Rosi, Peripheral T Cells as a Biomarker for Oxygen-Ion-Radiation-Induced Social Impairments. Radiat Res 190, 186–193 (2018).

