## Supplementary material for "Small molecule cognitive enhancer reverses age-related memory decline in mice": All supplemental figures

Supplemental Figure 1

A

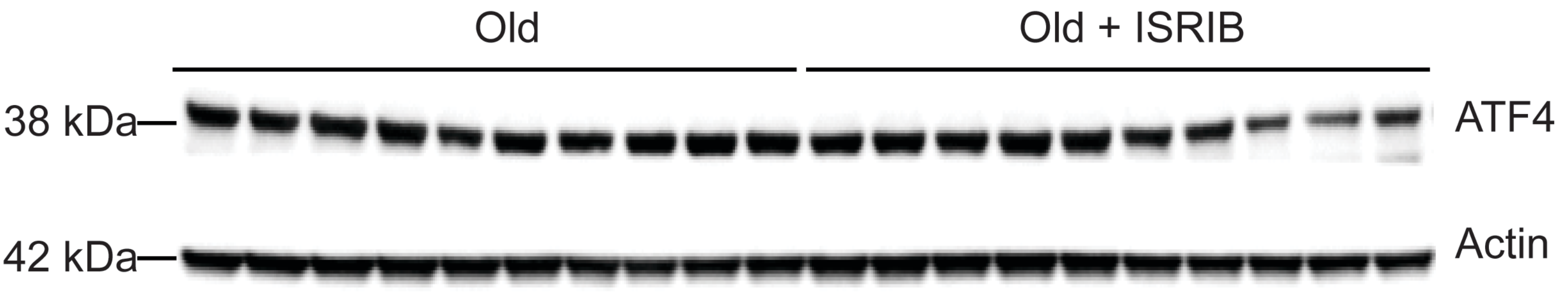


B

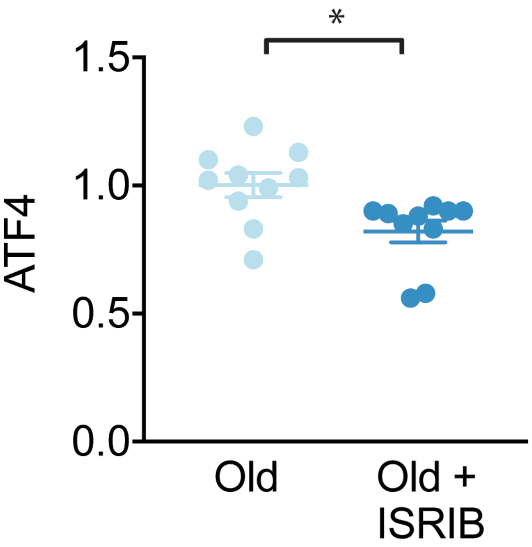

Supplemental Figure 2

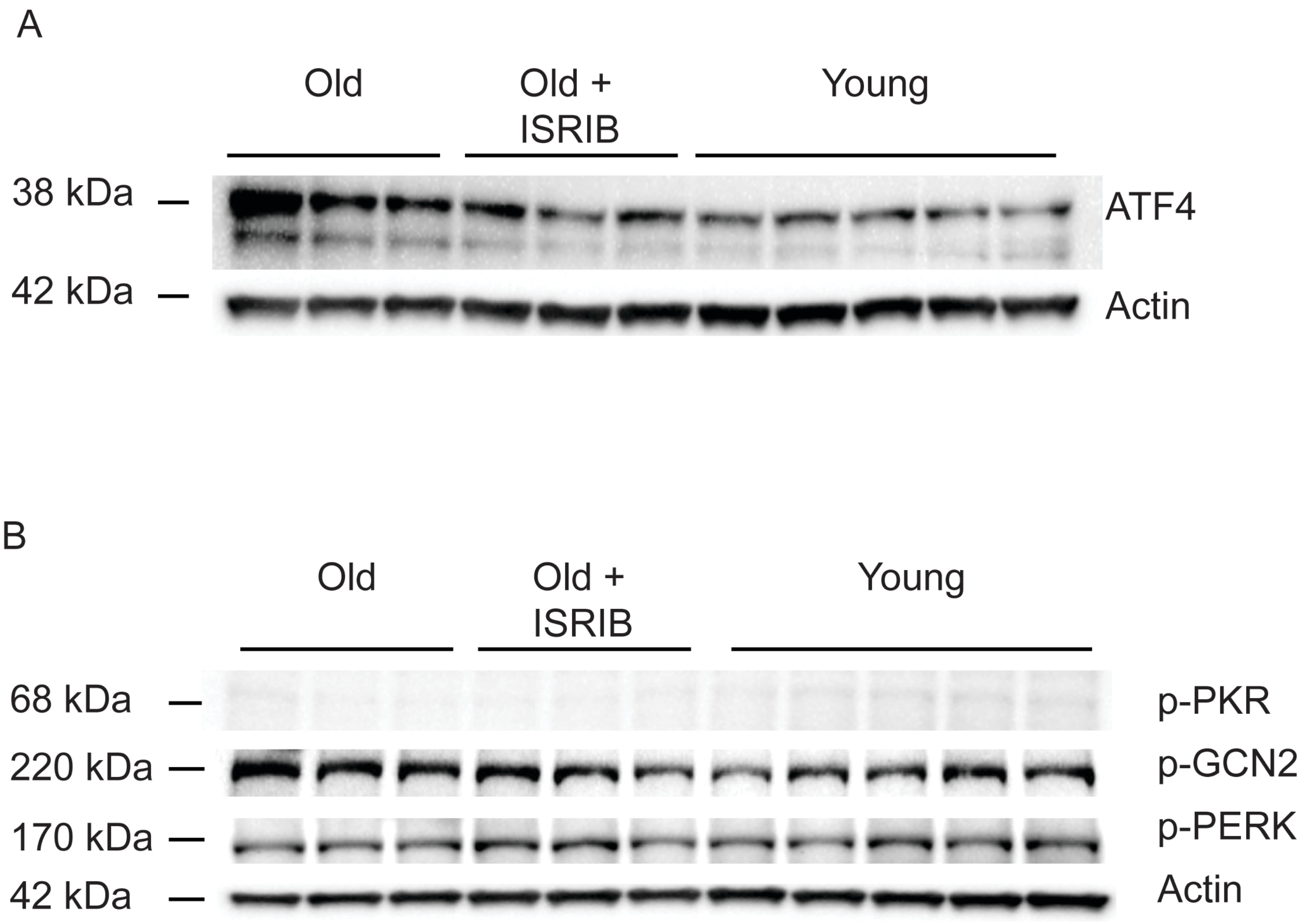

Supplemental Figure 3

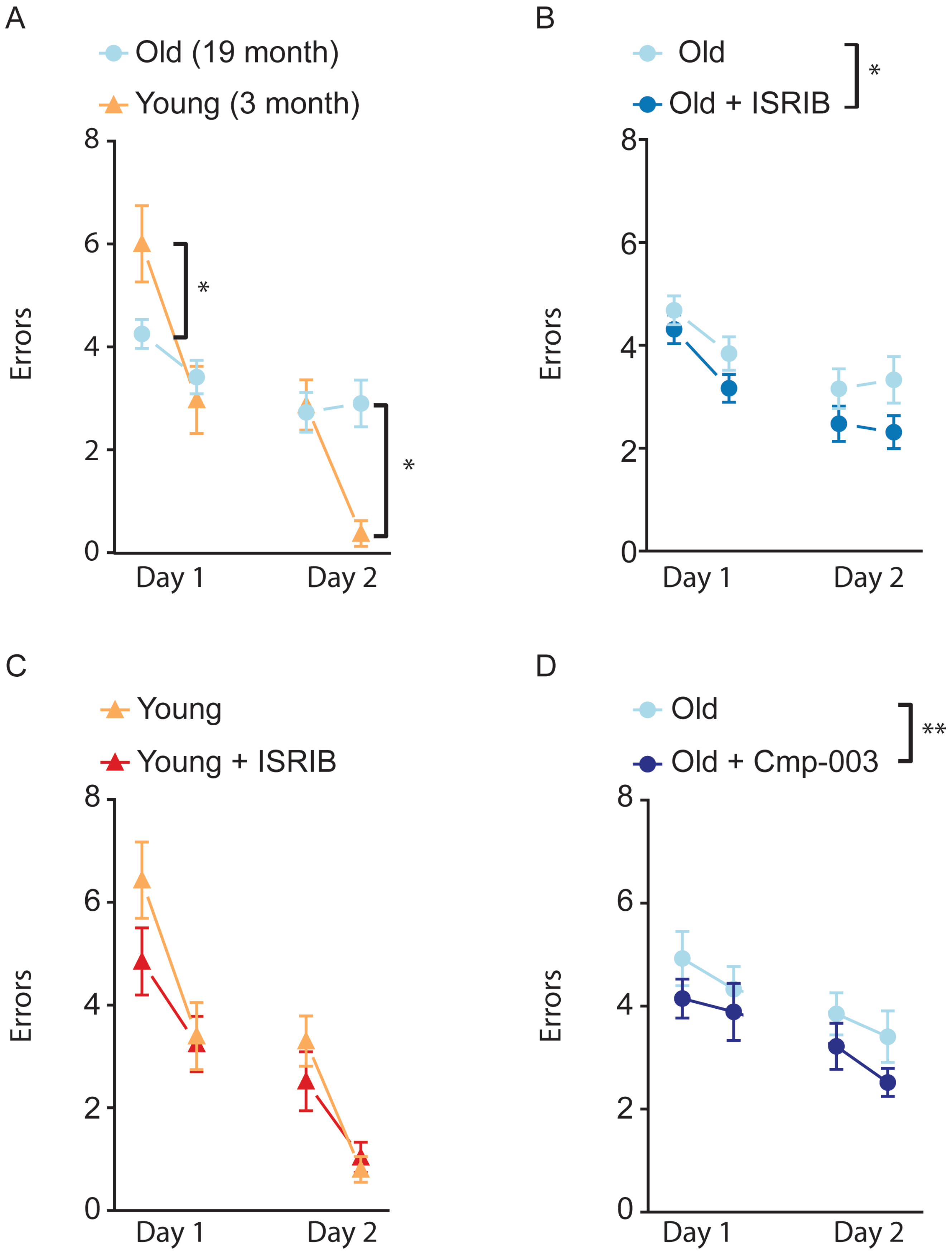

Supplemental Figure 4

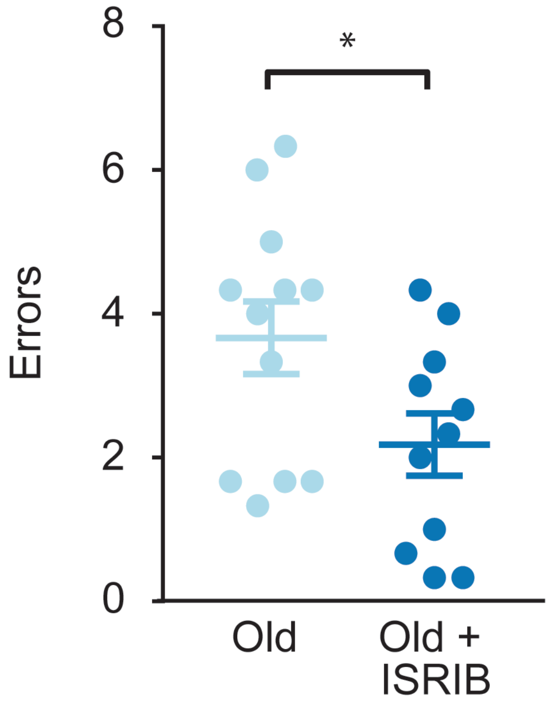

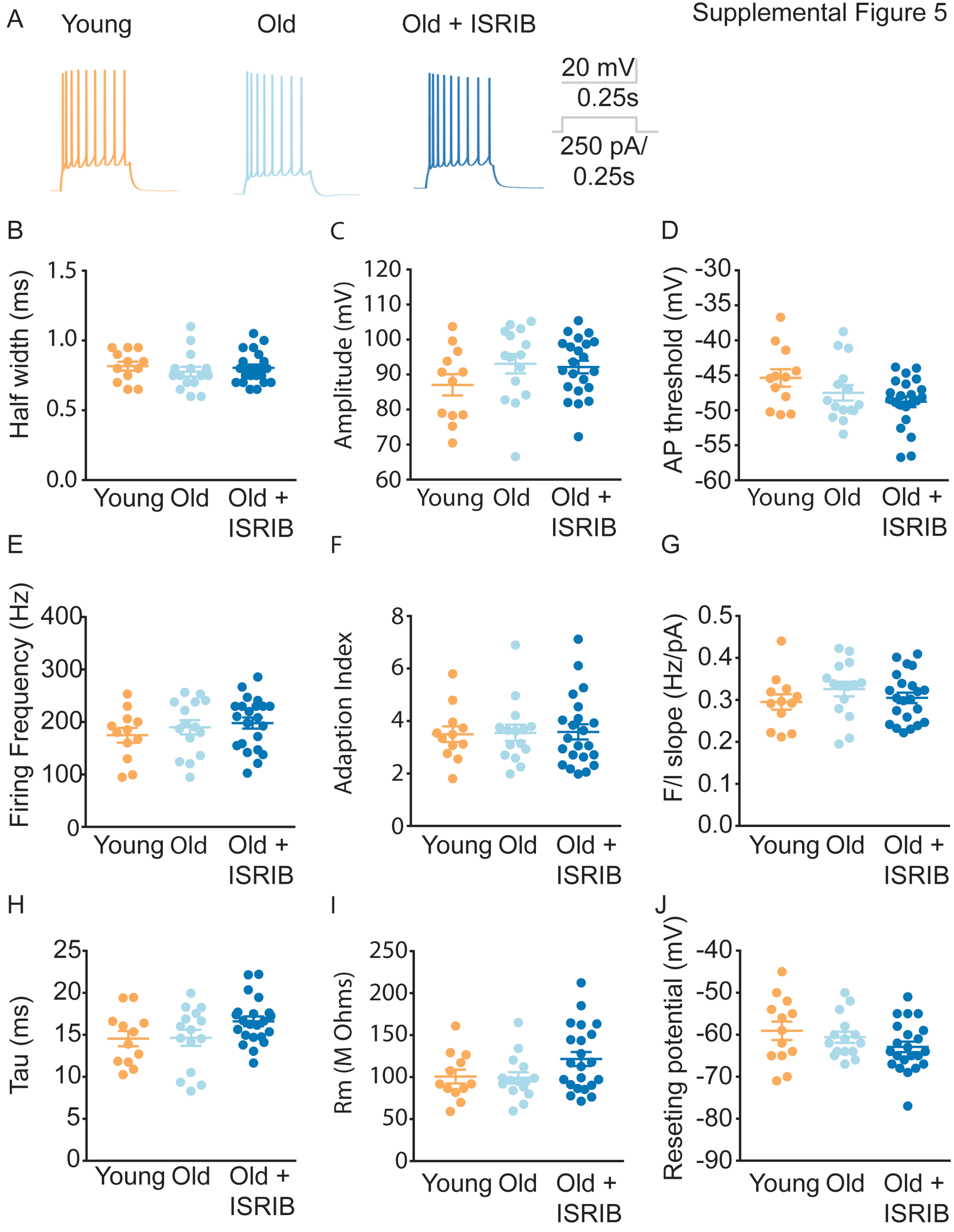
A
Young
Old
Old + ISRIB
20 mV
0.25s
250 pA/
0.25s
B
Half width (ms)
1.5
1.0
0.5
0.0
Young Old Old + ISRIB
C
Amplitude (mV)
120
110
100
90
80
70
60
Young Old Old + ISRIB
D
AP threshold (mV)
-30
-35
-40
-45
-50
-55
-60
Young Old Old + ISRIB
E
Firing Frequency (Hz)
400
300
200
100
0
Young Old Old + ISRIB
F
Adaption Index
8
6
4
2
0
Young Old Old + ISRIB
G
F/I slope (Hz/pA)
0.5
0.4
0.3
0.2
0.1
0.0
Young Old Old + ISRIB
H
Tau (ms)
25
20
15
10
5
0
Young Old Old + ISRIB
I
Rm (M Ohms)
250
200
150
100
50
0
Young Old Old + ISRIB
J
Reseting potential (mV)
-40
-50
-60
-70
-80
-90
Young Old Old + ISRIB

Supplemental Figure 6

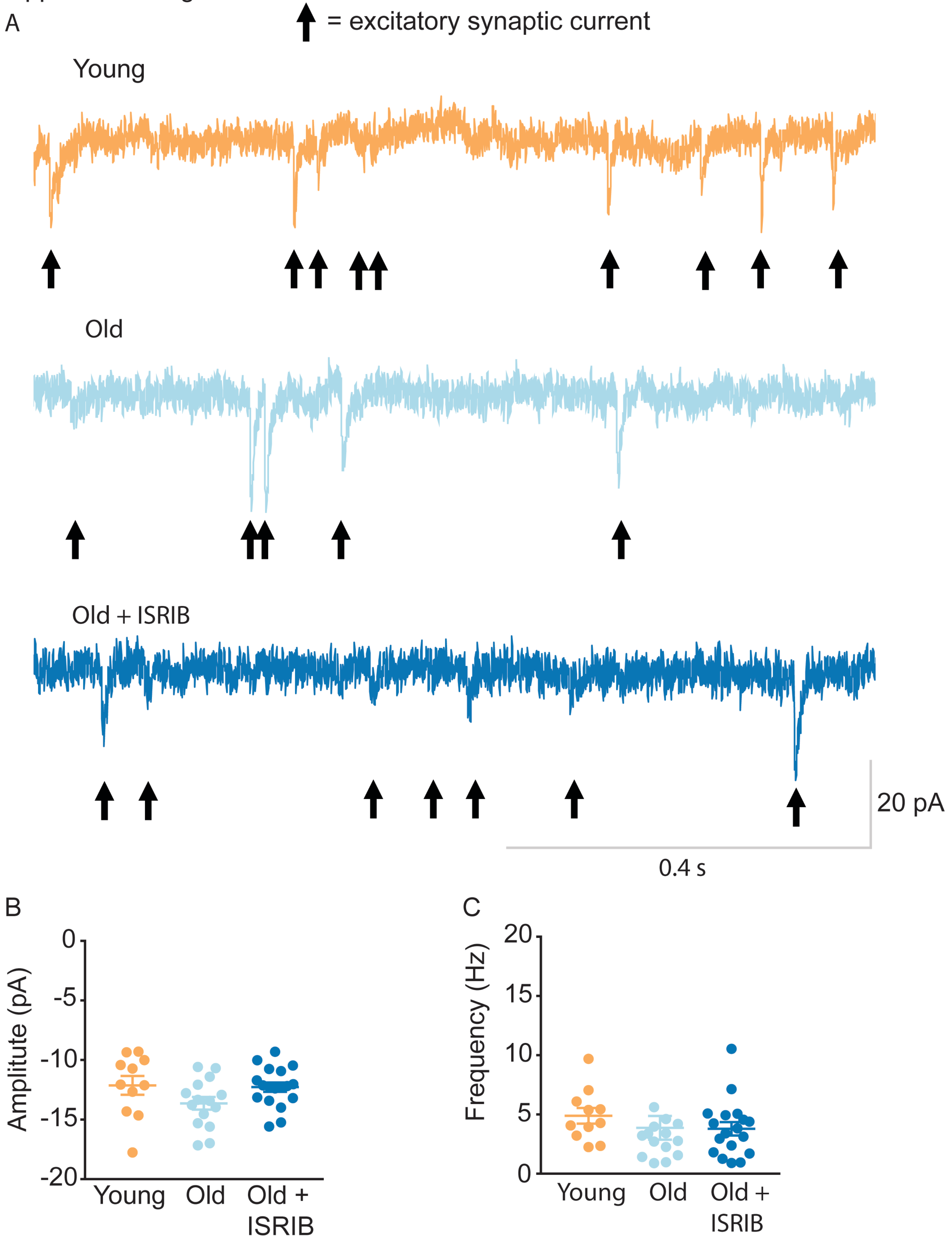

Supplemental Figure 7

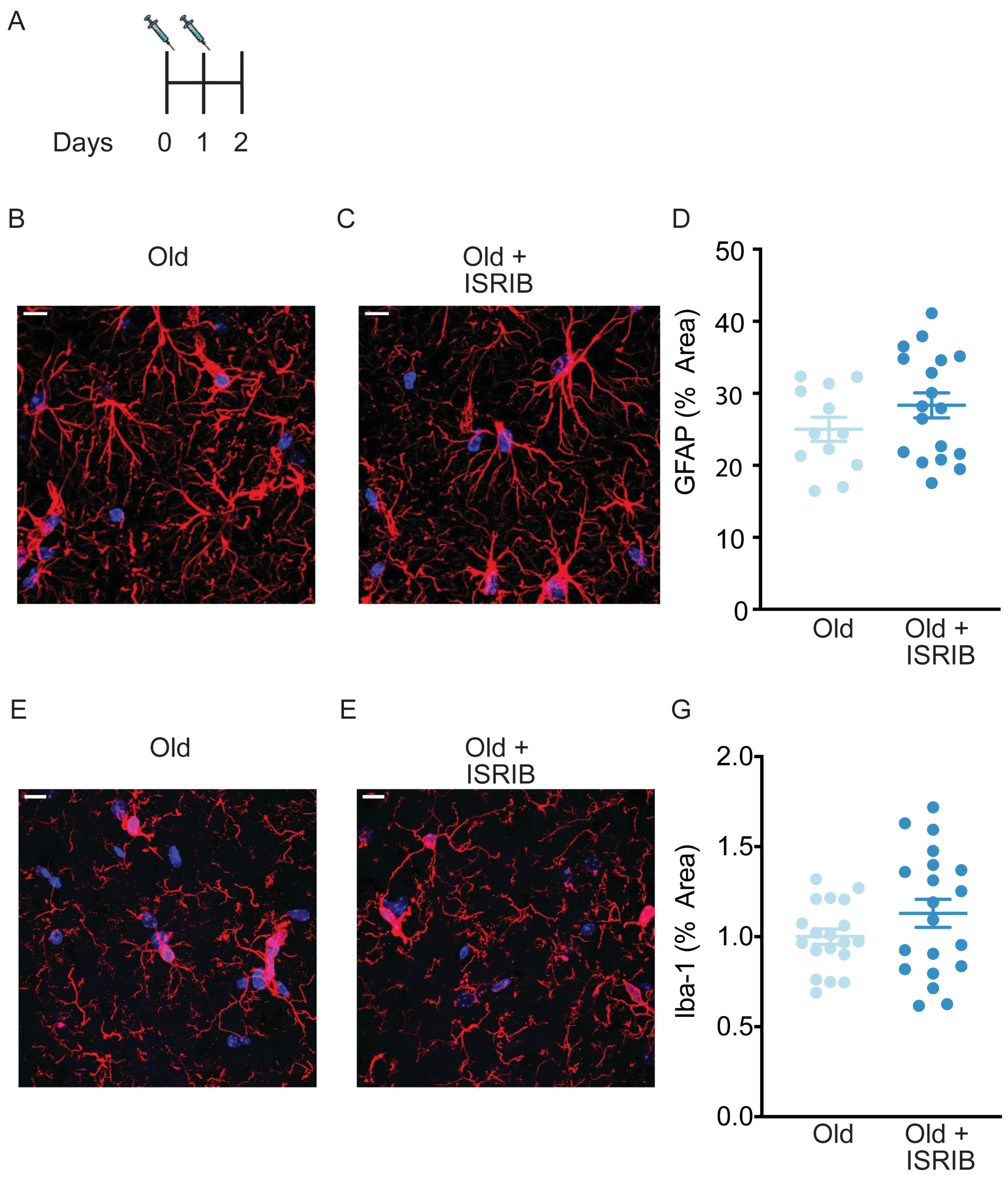

Supplemental Figure 8

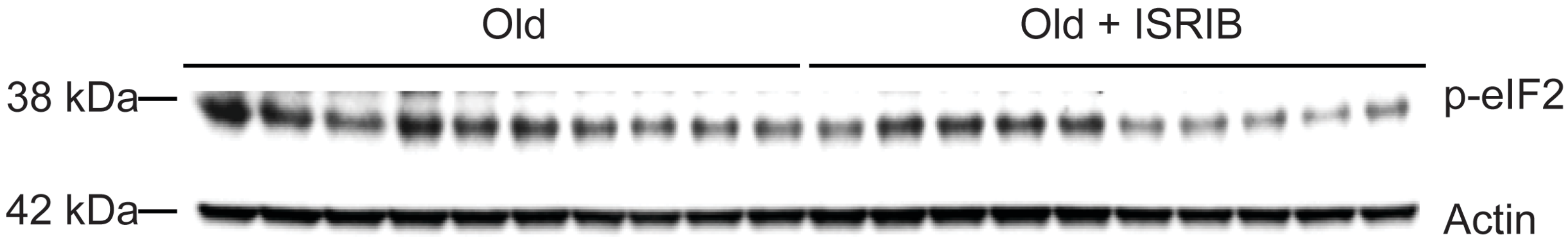
